# *RAS* mutations drive proliferative chronic myelomonocytic leukemia via activation of a novel KMT2A-PLK1 axis

**DOI:** 10.1101/2019.12.23.874487

**Authors:** Ryan M. Carr, Denis Vorobyev, Terra Lasho, David L. Marks, Ezequiel J. Tolosa, Alexis Vedder, Luciana L. Almada, Andrey Yurchenko, Ismael Padioleau, Bonnie Alver, Giacomo Coltro, Moritz Binder, Stephanie L. Safgren, Isaac Horn, Xiaona You, Nathalie Droin, Eric Solary, Maria E. Balasis, Kurt Berger, Christopher Pin, Thomas Witzig, Ajinkya Buradkar, Temeida Graf, Peter Valent, Abhishek A. Mangaonkar, Keith D. Robertson, Matthew T. Howard, Scott H. Kaufmann, Martin E. Fernandez-Zapico, Klaus Geissler, Eric Padron, Jing Zhang, Sergey Nikolaev, Mrinal M. Patnaik

**Affiliations:** Division of Hematology, Department of Internal Medicine, Mayo Clinic, MN, USA; INSERM U981, Gustave Roussy Cancer Center, Villejuif, France; Schulze Center for Novel Therapeutics, Division of Oncology Research, Mayo Clinic, MN, USA; McArdle Laboratory for Cancer Research, University of Wisconsin-Madison, Madison, WI, USA; INSERM U1170 and Department of Hematology, Gustave Roussy Cancer Center, Villejuif, France; Chemical Biology and Molecular Medicine Program, Moffitt Cancer Center, FL, USA; London Regional Transgenic and Gene Targeting Facility, Lawson Health Research Institute University of Western Ontario, London, Canada; 5TH Department of Internal Medicine I, Division of Hematology and Hemostaseology, Medical University of Vienna, Vienna, Austria; Ludwig Boltzmann Institute for Hematology and Hemostaseology, Medical University of Vienna, Vienna, Austria; Molecular Pharmacology and Experimental Therapeutics, Mayo Clinic, MN, USA; Department of Laboratory Medicine and Pathology, Mayo Clinic, MN, USA; Sigmund Freud University Vienna, Vienna, Austria

## Abstract

Chronic myelomonocytic leukemia (CMML) is an aggressive hematological malignancy with limited treatment options. Whole exome (WES) and targeted sequencing of several independent cohorts of CMML patients, comparing dysplastic (dCMML) to proliferative (pCMML) CMML, as well as paired chronic phase disease and acute leukemic transformation (LT), associate acquisition of oncogenic RAS pathway mutations, the most common being *NRAS^G12D^*, with aggressive disease and with disease progression. Using patient derived progenitor colony assays and a *NRAS*^G12D^-Vav-Cre mouse model, we further demonstrate the role of mutant RAS signaling in driving and maintaining pCMML phenotype. RNA-sequencing links RAS pathway mutations with an increased expression of genes encoding the mitotic checkpoint kinases PLK1 and WEE1. Further, we dmeoinstrated that non-mutated lysine methyltransferase KMT2A (MLL1) acts as mediator of NRAS-induced *PLK1* and *WEE1* expression. Finally, we demonstrate the translational value of our findings by showing that pharmacological PLK1 inhibition decreases monocytosis and hepatosplenomegaly while improving hematopoiesis in *RAS* mutant patient-derived xenografts. Hence, we define severe CMML as oncogenic RAS pathway-enriched malignancies, with a unique gene expression profile regulated by *KMT2A*, amenable to therapeutic intervention.

## INTRODUCTION

Chronic myelomonocytic leukemia (CMML) is an aggressive hematological malignancy characterized by sustained peripheral blood (PB) monocytosis (absolute monocyte count/AMC ≥1 x 10^9^/L), bone marrow (BM) dysplasia and an inherent risk for leukemic transformation (LT) to secondary acute myeloid leukemia (sAML) (Arber et al., 2016; Patnaik and Tefferi, 2018). The only curative modality, allogeneic hematopoietic cell transplantation (HCT), is rarely feasible because of age (median age at diagnosis 73 years) and frequent comorbidities (de Witte et al., 2017; Patnaik and Tefferi, 2018; Sharma et al., 2017). Patients that are not eligible for HCT receive symptom-guided therapies, ranging from erythropoiesis stimulating agents to hypomethylating agents (HMA) (Coston et al., 2019; Patnaik and Tefferi, 2018). While HMA epigenetically restore hematopoiesis in a subset of CMML patients, they fail to significantly alter the disease course, or impact mutational allele burdens in a vast majority; including responding patients, with patients even transforming to sAML while in a morphological complete remission (CR) (Coston et al., 2019; Merlevede et al., 2016). Finally, relative to patients with *de novo* AML, survival in blast transformed CMML (sAML) is much shorter (<4 months), even with the use of AML-directed therapies (Patnaik et al., 2018a). Thus there is an urgent and unmet need for rationally developed therapies for patients with CMML.

The 2016, iteration of the World Health Organization (WHO) classification of myeloid malignancies uses BM and PB blast percentages to subcategorize CMML into CMML-0, -1 and -2, with increasingly poor outcomes (Arber et al., 2016). The WHO also calls for the recognition of pCMML (proliferative) and dCMML (dysplastic) subtypes, based on a diagnostic white blood cell count (WBC) of ≥ 13 x 10^9^/L for the former (Arber et al., 2016). The molecular fingerprint for CMML combines recurrent mutations in approximately 40 genes, some of which are associated with poor outcomes (*ASXL1, NRAS, RUNX1* and *SETBP1*), and are incorporated into CMML-specific prognostic scoring systems (Elena et al., 2016; Itzykson et al., 2013a; Patnaik et al., 2014). Unresolved questions include how genetic and epigenetic events contribute to CMML progression and LT, and whether these molecular events render the CMML cells vulnerable to specific therapeutic interventions.

In the present study, we demonstrate that WHO-recognized disease severity criteria, *i.e.* cell proliferation and blast cell accumulation, correlate with acquisition of RAS pathway mutations, with *NRAS^G12D^* being the most common. Using next generation sequencing (NGS) studies, *in vitro* patient derived progenitor colony forming assays and a novel *Nras*^G12D^-Vav-Cre mouse model, we demonstrate that *NRAS* mutations drive the pCMML phenotype. In RAS mutant pCMML, the RAS pathway mutations result in increased expression of genes encoding the mitotic checkpoint kinases PLK1 and WEE1 through overexpression and displacement of the lysine methyltransferase KMT2A (MLL1), an enzyme that results in methylation of histone 3 (H3) lysine 4 (K4) at the promoters of *PLK1* and *WEE1*. While monomethylation of H3K4 is usually carried out by KMT2C/2D, we demonstrate that in RAS mutant pCMML this is specifically regulated by an unmutated and overexpressed KMT2A. We further show that pharmacological inhibition of PLK1 decreases monocytosis and hepatosplenomegaly and improves hematopoiesis in patient-derived, *RAS*-mutated CMML xenografts. These results emphasize the role of RAS pathway mutations in CMML progression, demonstrate for the first time an oncogenic function of a non-mutated but overexpressed KMT2A, and identify an opportunity for personalized therapeutic intervention in CMML.

## RESULTS

### RAS pathway mutations correlate with WHO-defined proliferative CMML (pCMML)

Analysis was first focused on chronic phase CMML, especially with regards to the phenotypic and survival differences between WHO-defined dCMML and pCMML subtypes. Univariate analysis of a cohort of 1183 WHO-defined Mayo Clinic-GFM-Austrian CMML patients (**Table S1**; dCMML, 607 and pCMML, 576), median age 72 years (range, 18-95 years), 66% male, with a median follow up of 50 months (range, 41-54.5 months) validated an inferior overall-survival (OS) of pCMML patients in comparison to dCMML (23 vs 39.6 months; p<0.0001) (**Figure 1A**). This stratification also predicted a shorter AML-free survival in patients with pCMML versus dCMML (**Figure 1B**, 18 vs 32 months; p<0.0001).

**Figure 1.**
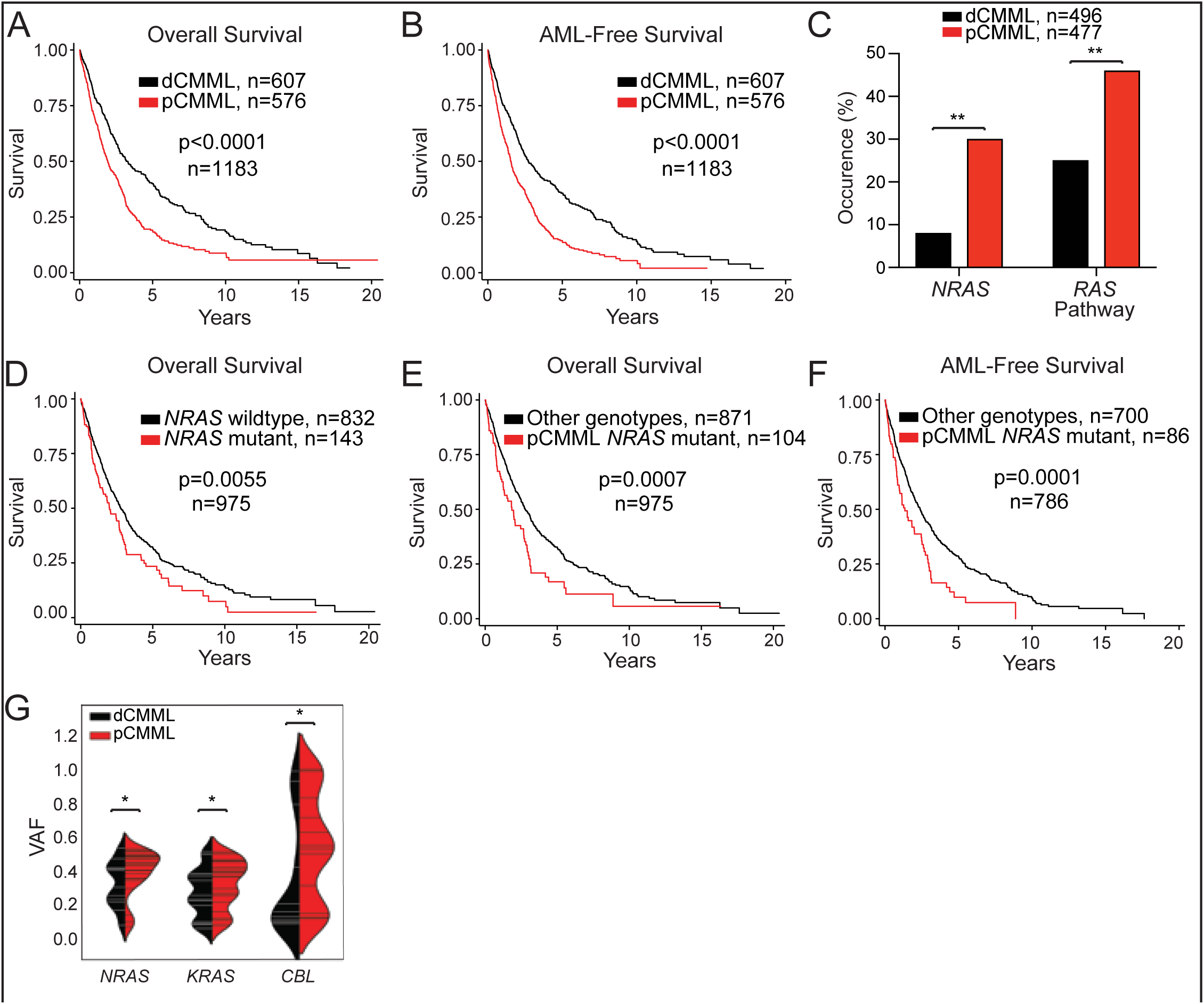
RAS pathway mutations correlate with WHO-defined proliferative chronic myelomonocytic leukemia (CMML). **A.** Kaplan-Meier curve depicting overall survival of CMML patients from the Mayo-GFM-Austrian cohort stratified by dCMML and pCMML subtypes. **B.** Kaplan-Meier curve depicting AML-free survival. **C.** Prevalence of *NRAS* and RAS pathway mutations in dCMML and pCMML by next generation sequencing. **D.** Kaplan-Meier curve depicting overall survival of CMML patients stratified by *NRAS* wildtype and *NRAS* mutant cases. **E.** Kaplan-Meier curve depicting overall survival in pCMML patients with *NRAS* mutations relative to other CMML genotypes. **F.** Kaplan-Meier curve depicting AML-free survival in pCMML patients with *NRAS* mutations relative to other CMML genotypes. **G.** Violin plots representing variant allele frequencies (VAFs) of the most frequent RAS pathway mutations in dCMML and pCMML. VAF is depicted on the y-axis. Width of horizontal hatches correlates to number of samples with the indicated VAF. * indicates p value < 0.05; ** indicates p value < 0.01. See also **Figure S1**.

Genetic differences were assessed between the two subtypes in 973 molecularly annotated Mayo Clinic-GFM-Austrian CMML patients, including 477 with pCMML and 496 with dCMML, using a targeted NGS assay that was designed to detect mutations in the following oncogenic RAS pathway genes; *NRAS, KRAS, CBL* and *PTPN11*. A higher frequency of *NRAS* (30% versus 8%; p<0.001) and cumulative oncogenic RAS pathway mutations (46% versus 25%, p<0.001) were identified in pCMML versus dCMML (**Figure 1C**; **Table S2**). Additional genetic differences between pCMML and dCMML also included a higher frequency of *ASXL1* (60% vs 42%; p<0.0001), *JAK2^V617F^* (12% vs 5%; p=0.00011) mutations and trisomy 8 (11% vs 4%; p=0.02) in pCMML, while *TET2* (43% vs 54%; p=0.002) mutations were more common in dCMML. There were no differences in the distribution of *SRSF2, SF3B1, U2AF1, IDH1, IDH2, DNMT3A, SETBP1, RUNX1, BCOR, TP53* and *PHF6* mutations between the two subtypes (**Table S2**).

Patients were then stratified by the presence or absence of *NRAS* mutations and demonstrated an inferior OS for *NRAS* mutant CMML in comparison to *NRAS* wildtype CMML (24 vs 33 months; p=0.0055) (**Figure 1D**). This impact was also retained for cumulative oncogenic RAS pathway mutations (**Figure S1A**). Given the predominance of *NRAS* mutations in pCMML, we further subdivided pCMML and dCMML into subsets with and without *NRAS* mutations. pCMML with *NRAS* mutations was the most aggressive CMML subtype, with significantly shorter OS (22 months; p=0.0007) relative to all other CMML subtypes combined (33 months) (**Figure 1E**). Notably, OS was not significantly different when comparing each CMML subtype stratified by *NRAS* mutational status (**Figure 1SB**). Similarly, *NRAS* mutant pCMML had the shortest AML-FS (16 months; p=0.0001) in comparison to other subclasses combined (AML-FS 29 months) (**Figure 1F**), and in comparison to pCMML without *NRAS* (AML-FS 19 months), dCMML with *NRAS* mutations (AML-FS 29 months) and dCMML without *NRAS* mutations (AML-FS 37 months) (**Figure S1C**). In addition, based on variant allele frequency (VAF) analysis, in 977 CMML patients that underwent NGS, there was a higher frequency of clonal *NRAS, KRAS* and *CBL* mutations in pCMML in comparison to dCMML (**Figure 1G, S1D**).

Paired WES performed on BM or PB mononuclear cells (MNC) collected at both chronic phase CMML and at AML-transformation stages in a cohort of 48 patients were then analyzed (**Table S3)**. Median OS was 25 months after diagnosis and 6.4 months after LT. At CMML diagnosis, or first referral to study institution, mutations in driver genes (detected in every patient; mean number, 3.6 per patient) and somatic copy number alterations (SCNA) (**Figure 2A-B, S2A**) were in accordance with previous reports (Merlevede et al., 2016). Some genetic alterations, such as mutations in *TET2*, *SRSF2* and *ASXL1*, were predominantly clonal while others, including those affecting genes of the oncogenic RAS pathway, such as *NRAS*, *KRAS* and *CBL*, showed a bimodal distribution of VAF (**Figure 2C**). Compared to chronic phase CMML, additional driver mutations (median = 2) were detected in 44% of LT samples (**Figure S2A**). The most striking changes included an increase in the prevalence of mutations in genes of the oncogenic RAS pathway (from 52 to 67%), and in SCNA (from 52 to 75%) (**Figure 2A-B, Figure S2A).**

**Figure 2.**
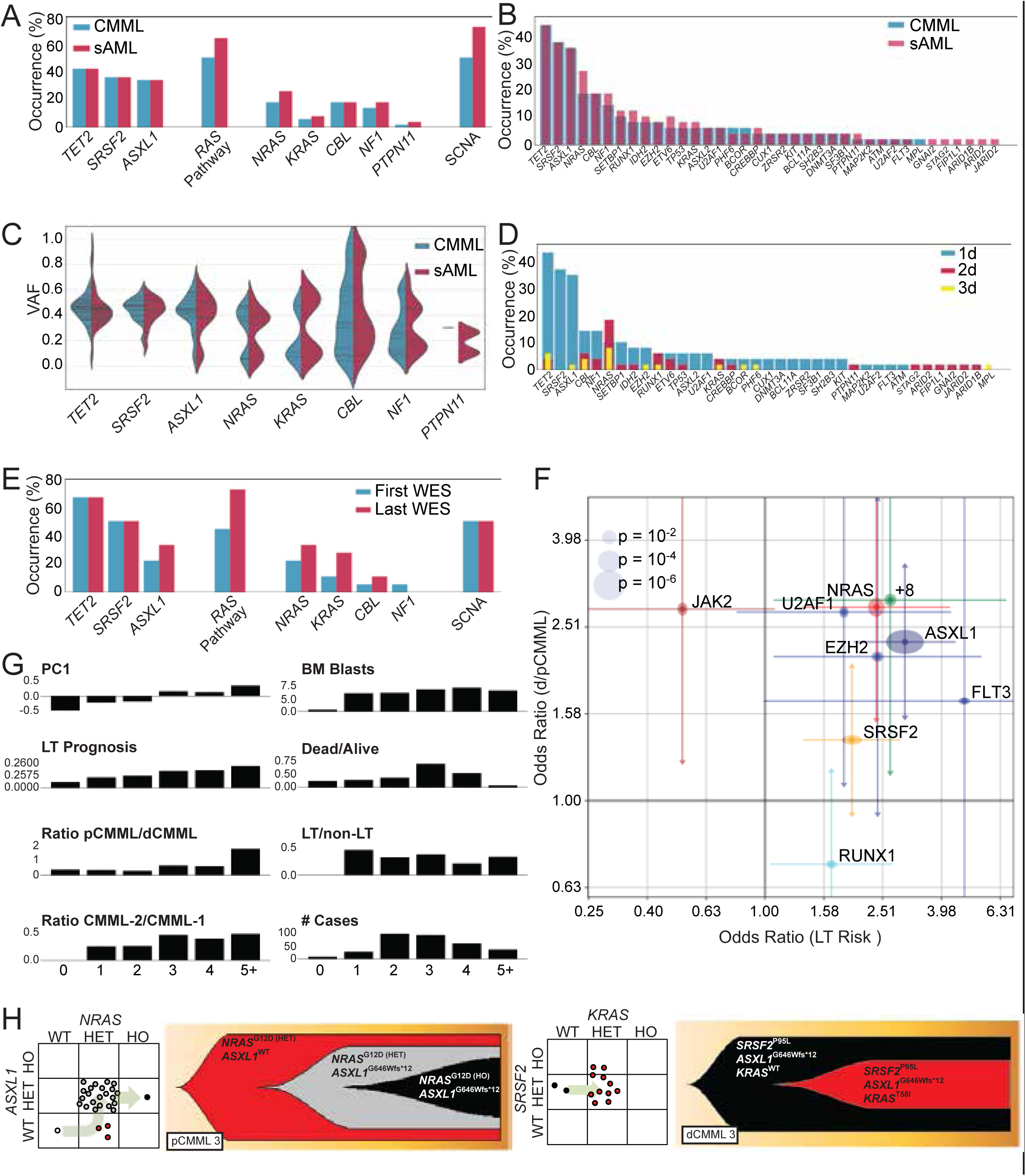
Alterations in driver mutation profiles delineate stages of CMML progression. Whole exome analysis of 48 paired CMML and secondary AML (sAML) samples. **A.** Prevalence of driver mutations and somatic copy number alterations (SCNAs) in CMML and secondary AML. RAS pathway represents all mutations in MAPK pathway. **B.** Frequency of driver mutations and SCNAs in CMML and sAML. **C.** Violin plots depicting variant allele frequencies (VAFs) for the most frequent driver mutations in CMML and sAML. **D.** Frequency of driver mutations and SCNAs in CMML and sAML and their categorization as 1d, 2d or 3d. **E.** Genetic profiling of CMML evolution in eighteen patients at two time points by whole exome sequencing. **F.** Odds ratios of genetic factors selected by the L1-regularized logistic regression that have the greatest impact on dysplastic/proliferative CMML categorization and leukemic transformation (LT) risk. The x-axis represents odds ratios of LT risk (high or low). The y-axis represents categorization of low or high white blood cell (WBC) count, or pCMML vs dCMML classification. The horizontal and vertical size of ellipses reflects corresponding p-values. Additionally, confidence intervals (alpha = 0.05) of odds ratio values are presented. **G.** Stratification of parameters associated with aggressive disease phenotype by number of driver mutations. **H.** Fish plots derived from patient-derived single colony assays of representative cases of pCMML (left) and dCMML (right). Scatter plot to the left of each fish plot represents the genotype of individual colonies as determined by Sanger sequencing of the indicated genes. *RAS* mutation status is depicted on the x-axis. The arrow indicates inferred evolutionary trajectory from which the fish plot was derived. See also **Figure S2**.

In order to trace clonal evolution from CMML onset to LT, we categorized driver mutations (Table S4 for definition of driver mutations) into those detected in both CMML and LT (referred to as primary drivers or 1d), those detected only in LT (secondary drivers, 2d) and those detected in CMML and lost in LT, suggesting sub clonal secondary drivers (3d) (**Figure S2B**). Sets of secondary drivers (2d and 3d) were strikingly different from primary drivers (1d), *i.e.* the average number of driver mutations per category (1d: 3.3; 2d: 0.8; 3d: 0.4) and the spectrum of mutated genes were clearly distinct. *TET2, ASXL1, SRSF2* mutations as well as monosomy 7 were almost exclusively 1d. *NRAS* was the main gene whose mutation rate increased from chronic phase to LT (**Figure 2D**). The fraction of *NRAS* mutations among driver mutations detected in LT was significantly higher than that detected in CMML at diagnosis (p=0.0002) (**Figure 2D, Figure S2C**). In 27% of cases, driver mutations were lost between chronic phase and LT. Eliminated sub clones harbored fewer driver mutations than progressive sub clones (19 vs 38; binomial test, p=0.01), suggesting that clones associated with LT underwent a longer evolutionary process than those eliminated with disease evolution (**Figure S2D**). CMML patients without oncogenic RAS pathway mutations in chronic phase had a high likelihood of acquiring them at LT (**Figure S2E**). In line with that, in the WES cohort, CMML patients without RAS pathway mutations had a better OS, in comparison to those with RAS pathway mutations (p<0.05).

WES of serial samples collected along disease evolution from 18 CMML patients (data set previously published) (Merlevede et al., 2016) showed the time-dependent accumulation of driver mutations that, again, affected mostly genes of the RAS pathway, including *NRAS*, *KRAS* and *CBL* (**Figure 2E**). In addition, In the combined WES cohort, white blood cell count (dCMML and pCMML subcategorization) as well as BM/PB blast fraction (CMML2/CMML1) also predicted LT risk (WBC [OR: 2.3, CI:1.4-3.9], blast fraction [OR: 5.1, CI:2.8-9.3]) and OS (WBC [OR: 1.7, CI:0.91-3.2]). Importantly, *NRAS* mutations were most significantly associated with aggressive disease features, such as leukocytosis (OR=3.0, CI: 1.7-5.1) and LT risk (OR=2.7, CI: 1.4-5.3), (**Figure 2F).**

Of note, the number of driver mutations correlated with the disease severity in CMML (**Figure 2G**). Targeted sequencing of CMML patient-derived single progenitor colony forming assays indicated that mutations can accumulate in multiple orders, e.g. mutations in *ASXL1* or *SRSF2* can appear after those in *NRAS* or *KRAS*, which prominently was the case in pCMML (**Figure 2H**). Altogether, these analyses in several independent patient cohorts indicate that enrichment in oncogenic RAS pathway mutations, with *NRAS* being most common, is typically associated with the pCMML phenotype, CMML related aggressive disease features and with CMML transformation to sAML.

### *NRAS* activation drives a proliferative CMML phenotype

To explore the impact of RAS pathway mutations on CMML cells, we used patient-derived cells to perform hematopoietic progenitor colony forming assays. Transduction of *NRAS* wild-type dCMML MNC with a wild-type *NRAS* expressing adenoviral vector construct resulted in a significant increase in colony formation, in comparison to empty vector controls (**Figure 3A**). *NRAS*^G12D^ transfection in *NRAS* wild-type CMML cells increased their *ex vivo* proliferation whereas siRNA depletion of *NRAS* in *NRAS*^G12D^ CMML cells demonstrated the opposite effect (**Figure 3B**).

**Figure 3.**
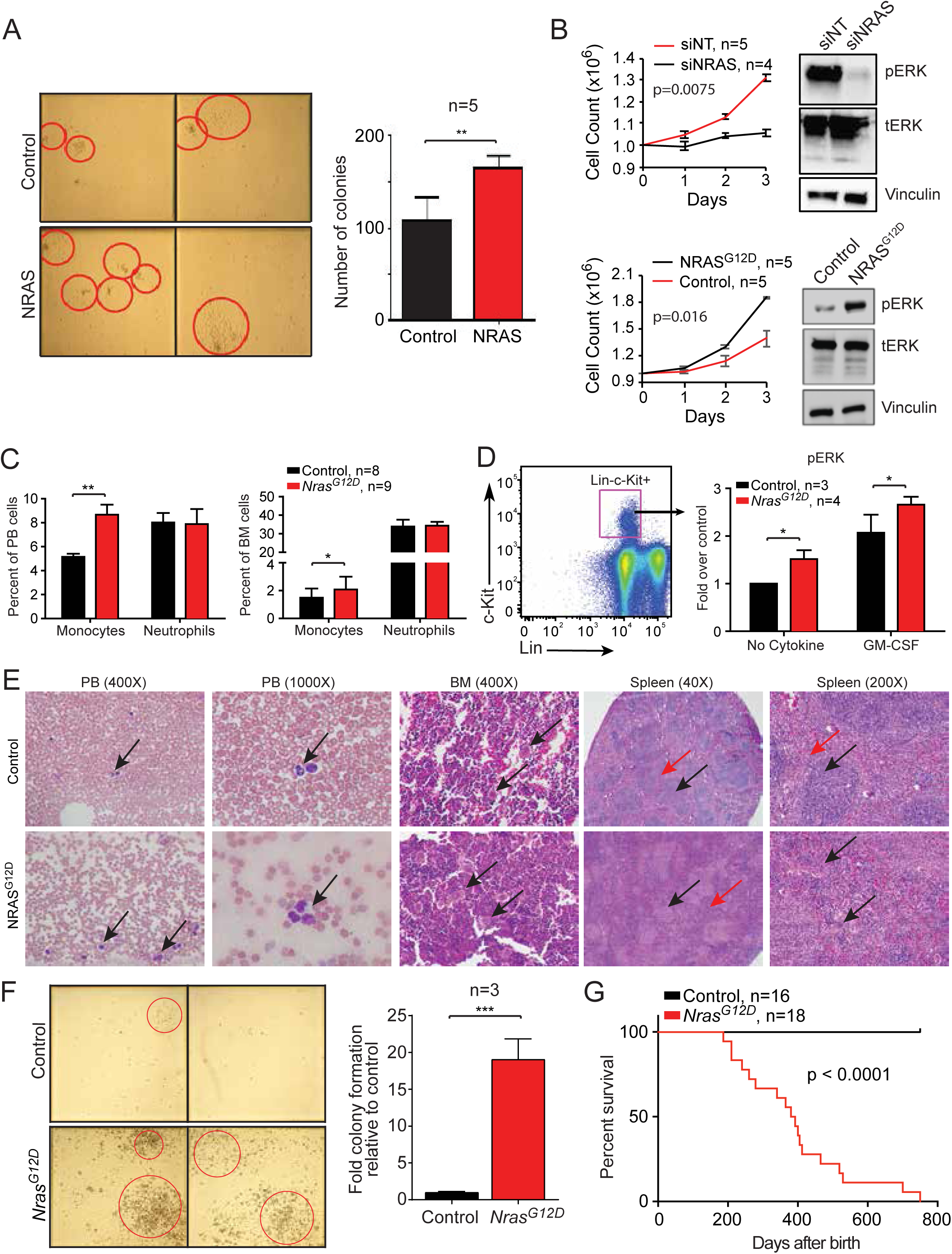
*NRAS^G12D^* mutation drives a proliferative CMML phenotype. **A.** Representative hematopoietic progenitor colony formation assay using *NRAS*-wildtype CMML patient-derived mononuclear cells (MNCs) after transduction with either a null adenoviral construct (Control) or NRAS-expressing vector (NRAS). Red circles indicate individual colonies. Bar graph depicts colony counts at day 12 after inoculation in five individual cases. **B.** Daily cell counts of CMML patient-derived MNCs after siRNA depletion of *NRAS* in *NRAS^G12D^*cells (above) and transfection of *NRAS^G12D^* in *NRAS* wildtype cells (below). Representative Western blots depicted for validation by assessing phosphorylated ERK (pERK) relative to total ERK (tERK) levels with Vinculin as a loading control. Knockdown of NRAS by qPCR was ≥ 85%. Characterization of the *Vav-cre-Nras^G12D^* mouse model of CMML is depicted at 6 weeks of age (panels C-E) and when moribund (panels F and G). **C.** Bar graphs depict differences in peripheral blood (PB, left) and bone marrow (BM, right) monocytes and neutrophils in *Nras^G12D^* mice relative to wildtype controls. **D.** BM cells were serum-and cytokine-starved for 2 hours at 37°C. Cells were then stimulated with or without 2 ng/ml of mGM-CSF for 10 minutes at 37°C. Levels of phosphorylated ERK1/2 (pERK) were measured using phosphor-flow cytometry. Myeloid progenitors are enriched in Lin^-/low^ c-Kit^+^ cells. Values are mean ± SE. **E.** Histopathologic comparisons between wildtype control (top) and *Nras^G12D^* (bottom) mouse PB, BM and spleen. PB smears depict monocytes (arrows). BM shows normal hematopoiesis (arrows, above) and megakaryocytic atypia and hyperplasia (arrows, below). Lower power spleen reveals normal white and red pulp (black and red arrows respectively, above) with effacement of this architecture (black and red arrows, below). High power spleen reveals normal white and red pulp architecture (black and red arrows respectively, above) and dysplastic megakaryocytes (arrows, below). **F.** Representative hematopoietic progenitor colony formation assay using MNCs from wildtype control (top) and *Nras^G12D^* (bottom) mice. Bar graph depicts colony counts at day 12 after inoculation. **G.** Kaplan-Meier curve demonstrating overall survival of *Nras^G12D^* mice relative to wildtype controls. ns is not significant. * indicates p value < 0.05; ** is p value <0.01; *** is p value < 0.001.

While RAS pathway mutations may be sub-clonal in some cases, as previously described, single colony assays in pCMML cases suggest it could serve as an early somatic event. To determine the effect of *Nras^G12D^*on hematopoiesis, a genetically engineered *Vav-Cre*-*Nras*^G12D^mouse model on a C57B1/6 background was developed (Joseph et al., 2013). We chose to develop this over the existing *Mx1-Cre-Nras*^G12D^ mouse model, where the mice develop a myeloproliferative disorder, but eventually die of a spectrum of hematological malignancies (Li et al., 2011). The *Mx1-Cre* transgene expresses Cre recombinase under the control of an inducible *Mx1* promotor, which is silent in healthy mice and active in the liver and hematopoietic cells after induction. Due to its activity in the liver, it often results in hepatic histiocytic infiltrates and the mice do not develop a CMML-specific phenotype. This is in contrast to our model, where the *Cre* recombinase under the control of the Vav promoter is more specifically active in the hematopoietic system and results in a disease that phenocopies pCMML. At 6 weeks, these mice demonstrated a significant increase in PB and BM monocyte counts, without significant changes in absolute neutrophil counts (**Figure 3C**). Analysis of ERK phosphorylation in their BM progenitor cells (Lin^-^,c-kit^+^) suggested an increased sensitivity to GM-CSF relative to controls, a previously described feature of human CMML (**Figure 3D**) (Padron et al., 2013). Histopathologic analyses showed BM hypercellularity with megakaryocytic atypia/dysplasia and splenic red pulp hyperplasia, architectural distortion, along with splenic extramedullary hematopoiesis (**Figure 3E**). BM MNC obtained from moribund *Vav-Cre-Nras^G12D^* animals generated an increased number of colonies, compared to control mice (**Figure 3F**). Finally, *Vav-Cre-Nras^G12D^* mice showed a significantly shortened survival relative to control mice (median 400 days vs not reached; **Figure 3G**). These data demonstrate that the *Nras*^G12D^ mutation alone expressed in *Vav*+ hematopoietic cells is sufficient to initiate CMML-like disease.

### RAS pathway mutations drive the enrichment in mitotic checkpoint kinases *PLK1* and *WEE1* in pCMML

To further assess the biological effects of RAS pathway mutations in CMML, bulk transcriptome sequencing (RNA-seq) was performed on PB MNC of a cohort of 25 CMML patients in chronic phase; 12 pCMML with RAS pathway mutations and 13 dCMML without RAS pathway mutations. RAS pathway mutated/pCMML samples demonstrated a unique gene expression profile with 3,729 up-regulated and 2,658 genes down-regulated genes in comparison to RAS pathway wildtype dCMML samples (**Figure 4A-B**). Unsupervised clustering analysis identified genes regulating mitotic and checkpoint processes as being preferentially expressed in RAS pathway-mutated pCMML. Pathway analysis (ingenuity) identified cascades preferentially expressed in RAS pathway-mutated pCMML as 1) mitotic roles of polo-like kinase (e.g. *WEE1, PLK1, PLK2*), 2) G2/M checkpoint regulation (e.g. *WEE1, AURKA, CHEK1*), 3) chromosomal replication (e.g. *CDC45, RPA3*), and 4) *BRCA1*-mediated DNA damage response (e.g. *PLK1*). Focusing on genes downstream of the oncogenic RAS pathway whose products could be targeted with currently available drugs (in clinical trials for myeloid neoplasms), we selected *PLK1*, which is a conserved kinase involved in the regulation of cell cycle progression and chromosome segregation, and *WEE1*, which is a nuclear kinase, regulating cell size by influencing CDK1 (Liu et al., 2016; Matheson et al., 2016). RT-qPCR analysis of *PLK1* and *WEE1* gene expression detected an overexpression of these two genes in 21 pCMML compared to 14 dCMML patient samples, agnostic to their RAS mutation status (**Figure 4C**).

**Figure 4.**
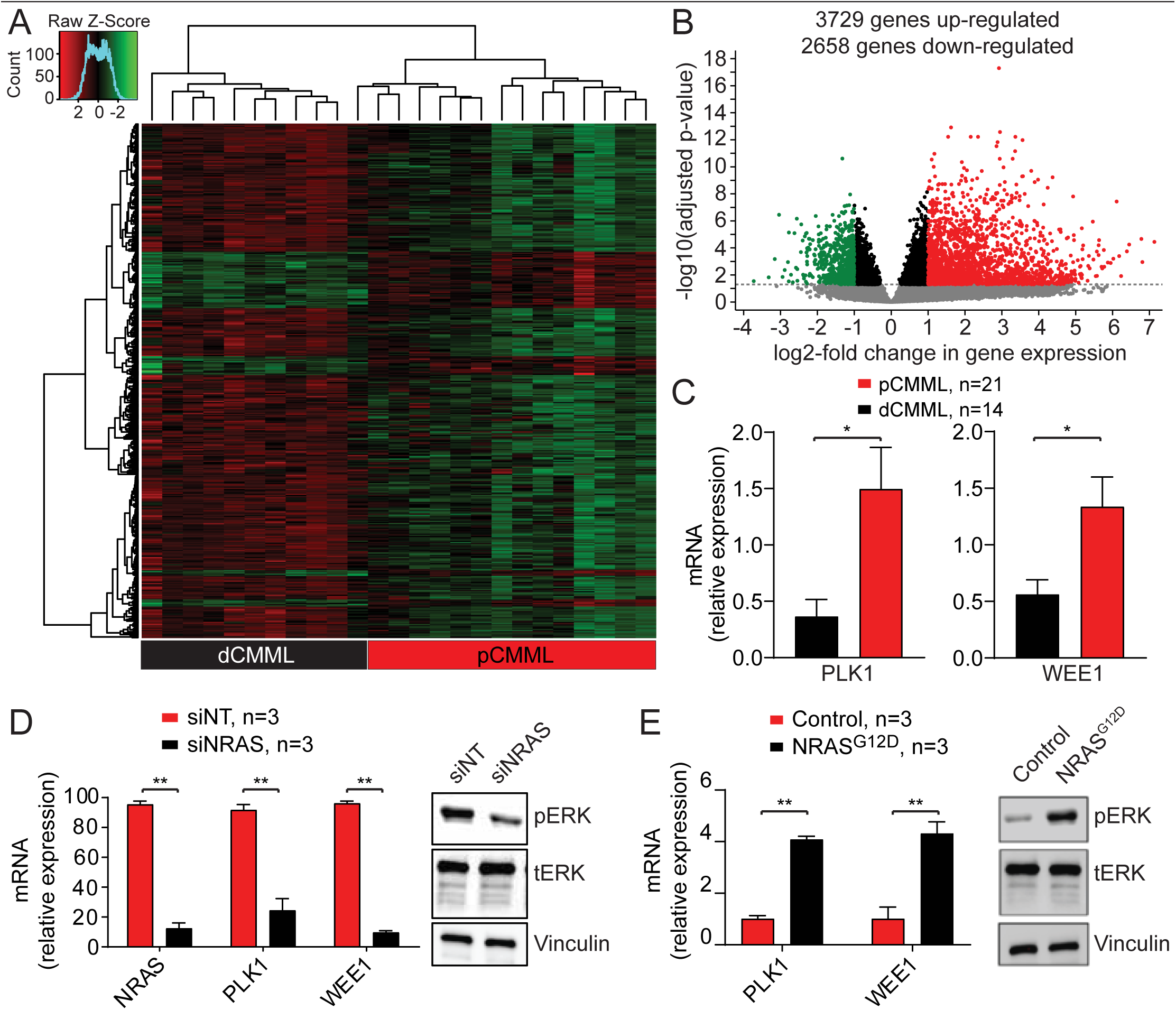
RAS pathway mutations drive expression of mitotic checkpoint kinases *PLK1* and *WEE1*. **A.** Unsupervised hierarchical clustering of RNA-seq performed on peripheral blood samples from 35 CMML patients. Cluster 1 (black) with dCMML cases and Cluster 2 (red) with pCMML cases. **B.** Volcano plot demonstrating differentially upregulated (red) and downregulated (green) genes in pCMML (vs dCMML) expressed as log2-fold change. **C.** Quantitative PCR (qPCR) validation of PLK1 (left) and WEE1 (right) expression in patient-derived MNCs from pCMML and dCMML cases. * indicates p value < 0.05. Data presented as mean ± SEM. **D.** qPCR for NRAS, PLK1 and WEE1 in NRAS mutant pCMML patient-derived MNCs after transfection with non-targeting siRNA (siNT) or siRNA against NRAS (siNRAS). Western blot is representative validation of NRAS knockdown. Data presented as mean ± SEM. **E.** qPCR for PLK1 and WEE1 in NRAS wildtype pCMML patient-derived MNCs after transfection with an empty vector (Control) or NRAS^G12D^. Western blot is representative validation of NRAS knockdown. Knockdown of NRAS by qPCR was ≥ 85%. Data presented as mean ± SEM. See also **Figure S4**.

Of note, this RNA-seq analysis was carried out on unsorted PB MNC after a pilot verification study that analyzed RNA-seq data from 5 CMML patients using unsorted PB MNC, 5 CMML patients using CD14+ sorted PB cells (monocytes) and 5 CMML patients using CD34+ selected PB progenitor cells (no overlap between groups), demonstrated a linear correlation in gene expression **(Figure S4A),** supporting the use of unsorted MNCs for further transcriptomic analysis in this study.

To further explore the role of RAS pathway mutations in *PLK1* and *WEE1* overexpression, we manipulated *NRAS* and performed RT-qPCR analyses. *NRAS* depletion by siRNA knockdown in *NRAS* mutant CMML cells resulted in a reduction in mitotic kinase expression, relative to controls (**Figure 4D**). Conversely, transfection of *NRAS*^G12D^ into *NRAS* wildtype CMML cells induced an upregulation of *PLK1* and *WEE1* expression, relative to the controls (**Figure 4E**). While the pCMML phenotype can also encountered in the context of *JAK2*^V61F^ (10%) (Patnaik et al., 2019), due to limited sample availability, we observed a trend towards higher *PLK1* and *WEE1* expression in *RAS* mutant/*JAK2* wildtype pCMML samples relative to *JAK2*^V61F^/ *RAS* wildtype pCMML. (**Figure S4B**). Together, these results suggest that acquisition of oncogenic RAS pathway mutations upregulate the mitotic check point kinases, *PLK1* and *WEE1,* in constituent pCMML cells.

### Mutant *RAS* regulates *PLK1* and *WEE1* expression through the lysine methyltransferase KMT2A (MLL1)

To explore the molecular events involved in *RAS*-mediated increase in *PLK1* and *WEE1* gene expression, we performed ChIP-seq (chromatin immunoprecipitation and NGS) experiments on PB and BM MNC collected from 40 CMML patients [RAS pathway mutated pCMML, *n*=18; 10 PB, 8 BM; RAS pathway wildtype dCMML, *n*=12, 8 PB, 4 BM) and healthy, age-matched controls (*n*=10, 2 PB, 8 BM). Among three studied histone marks, we noticed a global enrichment of monomethylated lysine 4 histone 3 (H3K4me1), a histone mark associated with gene activation, in pCMML relative to dCMML, without significant changes in H3K4me3 or H3K27me3 levels (**Figure 5A**, **Figure S5A**). On a sequence-specific analysis, an enrichment of H3K4me1 was detected at promoters of *PLK1* and *WEE1* (**Figure 5B**). *NRAS* siRNA-mediated knockdown decreased H3K4me1 occupancy at the promoter sites of *PLK1* and *WEE1* in RAS mutant CMML patient-derived MNCs relative to controls (**Figure 5C**). Conversely, *NRAS*^G12D^ transfection of *NRAS* wildtype patient-derived CMML cells resulted in an increased occupancy of H3K4me1 at the promoter sites of *PLK1* and *WEE1* (**Figure 5D**). In contrast, depletion of *JAK2*^V617F^ had no significant effect on H3K4me1 occupancy at the *PLK1* and *WEE1* promoters (**Figure S5B**) underscoring the specificity of this axis. Thus, oncogenic RAS pathway signaling likely enhances the expression of these mitotic kinases through enrichment of H3K4me1 at their respective gene promoters. Conversely, we did not see any difference in the occupancy of H3K4me3 and H3K27me3, between RAS mutant pCMML and RAS wild type dCMML at the promotors of *PLK1* and *WEE1* (**Figure S5A**).

**Figure 5.**
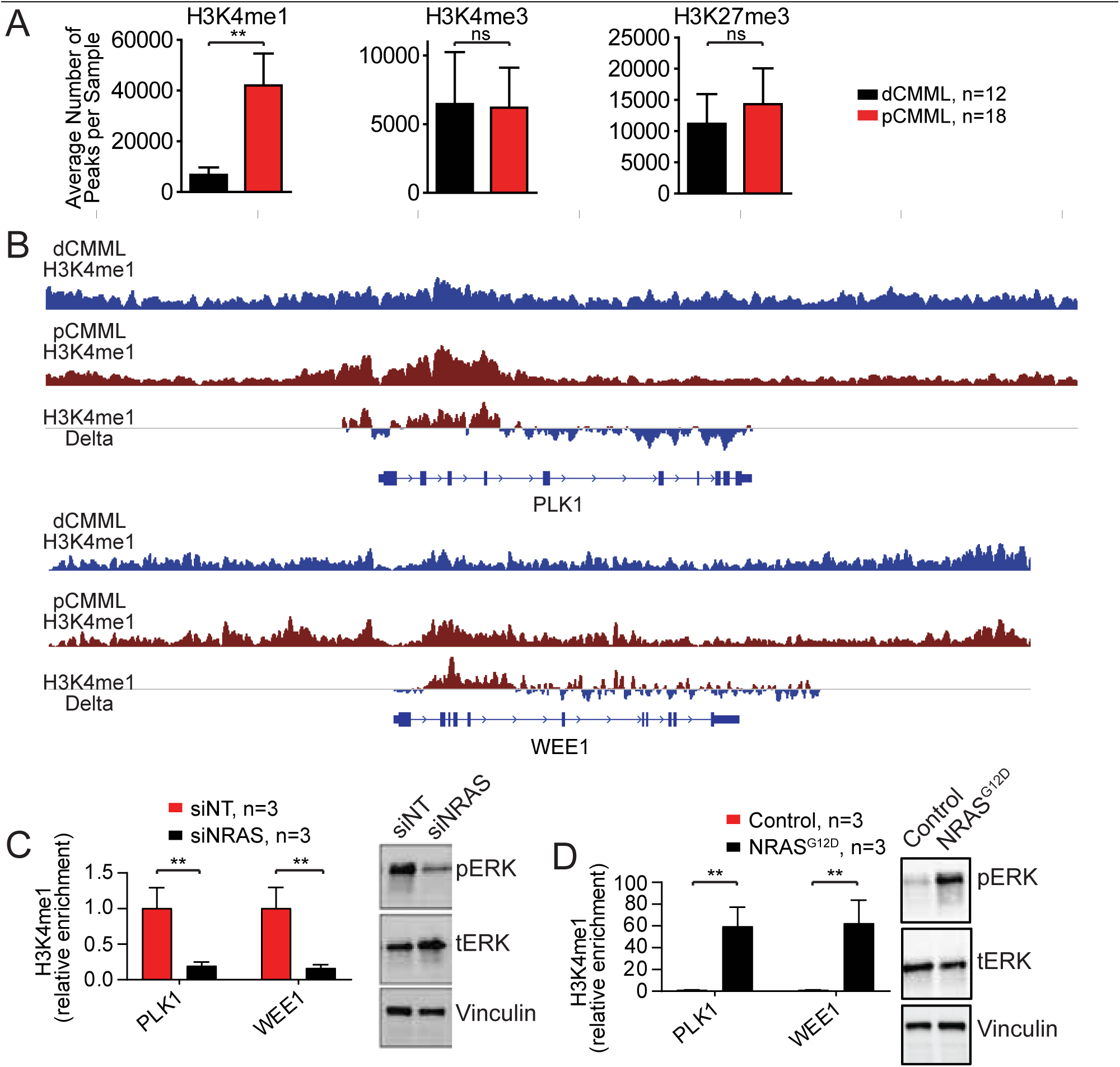
Genome-wide and sequence specific elevation of the H3K4me1 histone mark in pCMML. **A.** ChIP-seq on PB MNCs from CMML patients assessing relative global enrichment of histone 3 lysine 4 monomethylation (H3K4me1), trimetylation (H3K4me3), and histone 3 lysine 27 trimethylation (H3K27me3). Data presented as mean ± SEM. **B.** Sequence-specific ChIP-seq assessing H3K4me1 occupancy at *PLK1* (above) and *WEE1* (below) loci. dCMML (blue) and pCMML (red) traces indicate average signals derived from 12 and 18 patient samples respectively. The third trace represents a subtraction of these signals to assess differences. Relative localization along the *PLK1* and *WEE1* gene bodies is depicted. **C.** ChIP-PCR of *NRAS* mutant CMML patient-derived MNCs assessing *PLK1* and *WEE1* promoter occupancy of H3K4me1 after transfection with siNT or siNRAS. Data presented as mean ± SEM. Western blot is representative validation of NRAS knockdown. **D.** ChIP-PCR of *NRAS* wildtype CMML patient-derived MNCs assessing *PLK1* and *WEE1* promoter occupancy of H3K4me1 after transfection with a Control vector of NRAS^G12D^. Western blot is representative validation of NRAS knockdown. Knockdown of NRAS by qPCR was ≥ 85%. ** indicates p value < 0.001; ns is not significant. Data in panels A, B and D are presented as mean ± SEM. See also **Figure S5**.

To address the mutational impact of additional epigenetic regulator genes, especially the frequent occurrence of *TET2* mutations on the pCMML phenotype, we first performed DIP-seq (DNA immunoprecipitation and NGS) to assess changes in 5-methylcytosine (5-mc) and 5-hydroxymethylcytosine (5-hmc) in a sequence specific manner. We sequenced DNA extracted from BM-derived MNC in 18 patients, 9 with RAS pathway mutant pCMML (77% with *TET2* mutations) and 9 with RAS pathway wildtype dCMML (55% with *TET2* mutations) and found no differences in 5-mc and 5-hmc levels at the transcription start sites of *PLK1*and *WEE1* (**Figure S5C**, data not shown) between pCMML and dCMML. Thus, while *TET2* certainly has a role in CMML pathogenesis and loss of *TET2* has been shown to cooperate with *NRAS*^G12D^ in developing CMML (Kunimoto et al., 2018), oncogenic RAS signaling clearly remains the driver event influencing the pCMML phenotype.

Since monomethylation of H3K4 is carried out by the lysine methyltransferases KMT2A-D, SET1A and SET1B (Kerimoglu et al., 2017), we focused our subsequent work on these entities. RNA-seq analysis detected the overexpression of *KMT2A* (*MLL1*) and its coactivator *MEN1* in RAS mutant pCMML samples relative to RAS wild-type dCMML patients and healthy volunteers (**Figure 6A**). Interestingly, unlike in AML where *KMT2A* fusions and partial tandem duplications (PTD) are frequent, FISH (fluorescence in situ hybridization) and WES analyses in 500 CMML samples identified only a single patient with a *MLLT3-KMT2A* fusion (**Figure 6B**), without any *KMT2A* mutations or PTD (Dohner et al., 2017; Patnaik et al., 2018b), suggesting that overexpression of the wildtype gene could drive oncogenic activity of KMT2A in *RAS* mutant CMML. ChIP-PCR studies in MNC obtained from *NRAS* mutant pCMML patients detected occupancy of both KMT2A and MEN1 at the promoters of *PLK1* and *WEE1* (**Figure 6C and Figure S6A-B**). Knockdown of *NRAS* by siRNA in *NRAS* mutant pCMML MNC decreased occupancy of KMT2A at the promoter sites of *PLK1* and *WEE1* in comparison to siNT controls (**Figure 6D**). Conversely, transfection of *NRAS* wildtype dCMML patient samples with an *NRAS^G12D^* construct resulted in enhanced KMT2A occupancy at these aforementioned mitotic kinase promoters (**Figure 6E**). Further, siRNA-mediated knockdown of *KMT2A* conferred decreased H3K4me1 enrichment at the *PLK1* and *WEE1* promotor sites and decreased the mRNA expression of these mitotic kinase genes (**Figure 6F**). In addition, inhibition of the KMT2A-MEN1 interaction with Mi-503, a novel, orally bioavailable small molecule inhibitor (Borkin et al., 2015), similarly demonstrated decreased enrichment of H3K4me1 at *PLK1* and *WEE1* promoter sites (**Figure S6C**). Quantitative PCR studies at day 4 after MI-503 treatment confirmed decreased expression of *PLK1* and *WEE1* (**Figure S6D**). Since KMT2C and KMT2D are well known to monomethylate H3K4, we knocked down *KMT2C* and *KMT2D* using siRNAs in *NRAS* mutant pCMML MNC and demonstrated no changes in the enrichment of H3K4me1 at the *PLK1* and *WEE1* promotor sites, in comparison to siNT controls (**Figure S6E**). Together, these data support the hypothesis that *PLK1* and *WEE1* gene expression in *RAS* mutant CMML is dependent on an unmutated and overexpressed *KMT2A* as a mediator of oncogenic RAS signaling.

**Figure 6.**
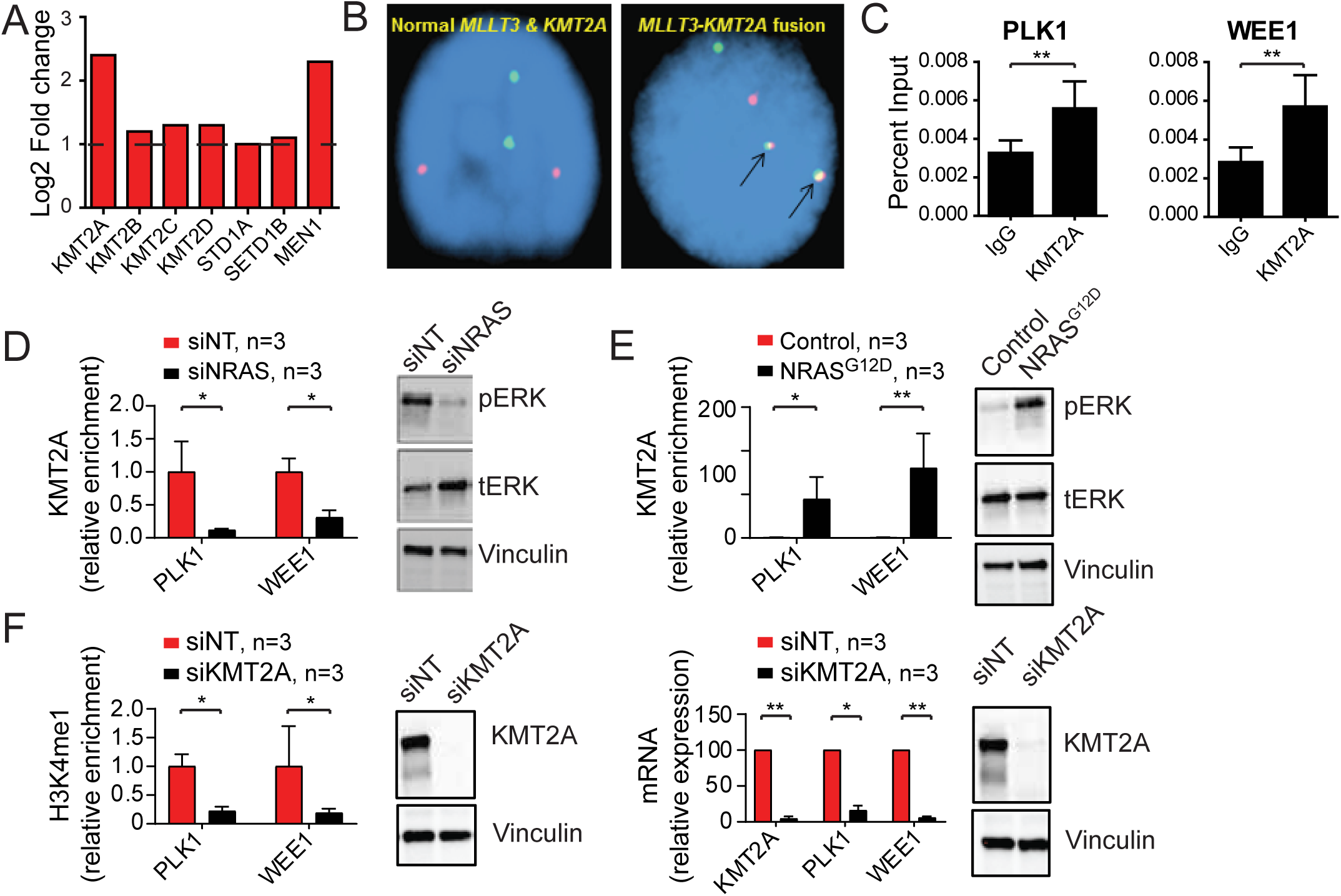
Mutant *RAS* regulates PLK1 and WEE1 expression through the lysine methyltransferase KMT2A (MLL1). **A.** RNA-seq data of expression levels of enzymes potentially involved in monomethylation in pCMML (vs dCMML and healthy volunteers). **B.** Representative fluorescence in situ hybridization (FISH) in two cases assessing for *MLLT3-KMT2A* fusion (n=500). Arrows (right) indicate presence of fusion. **C.** ChIP-PCR in *NRAS* mutant CMML patient-derived MNCs assessing KMT2A occupancy at promoters of *PLK1* (left) and *WEE1* (right) with immunoprecipitation of KMT2A relative to isotype control (IgG). **D.** ChIP-PCR assessing KMT2A occupancy at *PLK1* and *WEE1* promoters in *NRAS* mutant CMML patient-derived MNCs after transfection with siNT or siNRAS. Western blot is representative validation of NRAS knockdown. **E.** ChIP-PCR assessing KMT2A in *NRAS* wildtype CMML patient-derived MNCs after transfection with Control vector or NRAS^G12D^. Western blot is representative validation of NRAS knockdown. Knockdown of NRAS by qPCR was ≥ 85%. **F.** ChIP-PCR assessing H3K4me1 at promoters of *PLK1* and *WEE1* (left) and qPCR assessing levels of KMT2A, PLK1 and WEE1 in *NRAS* mutant CMML patient-derived MNCs after transfection with siNT or siKMT2A. Western blots are representative validation of KMT2A knockdown. Knockdown of KMT2A by qPCR was ≥ 90%. * indicates p value < 0.05; ** indicates p < 0.01. Data in panels C-F are presented as mean ± SEM. See also **Figure S6**.

### Therapeutic efficacy of targeting PLK1 in pCMML

Individual depletion of the mitotic kinases *PLK1* and *WEE1* using siRNA in patient-derived *RAS* mutant CMML cells resulted in reduced cell growth relative to siNT controls (**Figure 7A**). PLK1 inhibition with volasertib (Matheson et al., 2016) and WEE1 inhibition with AZD-1775 (Matheson et al., 2016; Ottmann et al., 2018) also efficiently reduced the generation of progenitor colonies derived from *NRAS* mutant pCMML samples, in comparison to *NRAS* wildtype dCMML (*N*=10) controls (**Figure 7B**). The IC_50_ levels for volasertib were 29.9 nM in *NRAS* mutant pCMML and 0.14 M for *NRAS* wildtype dCMML, while the values for AZD-1775 were 53.6 nM in *NRAS* mutant pCMML and 346 nM in *NRAS* wildtype dCMML.

**Figure 7.**
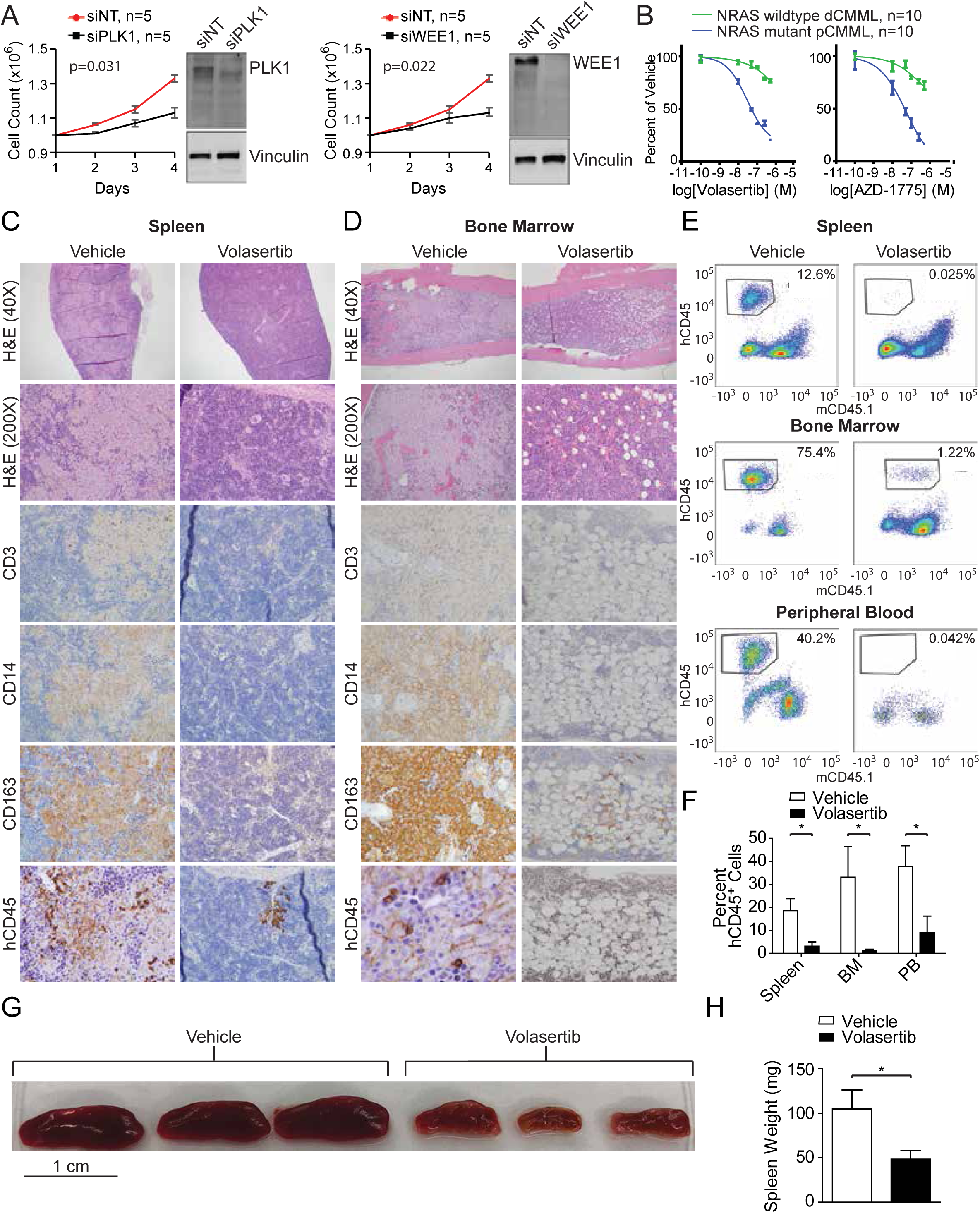
Therapeutic efficacy of targeting PLK1 in pCMML. **A.** Daily cell counts of CMML patient-derived MNCs after transfection with either siNT, siPLK1 (left) or siWEE1 (right). Representative Western blots depict validation of PLK1 and WEE1 knockdown. Knockdown of PLK1 and WEE1 by qPCR was ≥ 90%. **B.** Progenitor colony forming assays using CMML patient-derived MNCs with increasing doses of Volasertib (left) and AZD-1775 (right). Data in panels A and B are presented as mean ± SEM. **C and D.** Histopathologic analysis with H&E staining and immunohistochemistry (IHC) of spleen (C) and BM (D) of murine patient-derived xenografts (PDX) after treatment with vehicle control or Volasertib. Magnification is 200X unless otherwise indicated. hCD45 is human CD45. **E.** Representative flow cytometry of spleen, BM and PB of PDXs after treatment with vehicle control (left) or Volasertib (right). The y-axis indicates hCD45 expression status while x-axis indicates murine CD45 (mCD45). Percentages indicate proportion of hCD45+ and mCD45-cells in the respective tissues. **F.** Flow cytometry of proportion of hCD45+ cells in spleen, BM and PB of PDX mice. Data are presented as mean ± SEM from 9 mice. **G and H.** Effect of vehicle versus Volasertib treatment on PDX spleen size (G) and weight (H). * indicates p value < 0.05. Data are presented as mean ± SEM from 9 mice. See also **Figure S7**.

Finally, *NRAS*^G12D^ mutant pCMML-patient derived xenografts (PDX; NSG-SGM3 mice) (Yoshimi et al., 2017) were treated with volasertib. Six-week-old NSG-SGM3 mice were sub-lethally radiated with 25 Gy of radiation and then, 18 hours later, were injected with 2 x10^6^ CMML patient-derived BM MNCs via their tail veins (**Figure S7**). Four weeks later, after confirming engraftment by documenting hCD45 by flow cytometry, triplicate sets of mice were treated with volasertib 20 mg/kg/week administered intraperitoneally for 2 weeks versus a vehicle only control. After 2 weeks of therapy, the volasertib and vehicle-treated mice were sacrificed, followed by extensive flow cytometry-based and histopathological analysis of blood, BM and spleen specimens (**Figures S7**). These experiments demonstrated a marked reduction in BM and splenic malignant monocytic/histiocytic tumor infiltrates in volasertib-treated mice relative to vehicle-treated controls (**Figures 7C-D**). Immunohistochemistry indicated that neoplastic cells expressed hCD45, CD14 and CD163 while not expressing hCD3 and hCD19 (**Figure 7C-D and not shown for hCD19**). In addition, there was a partial normalization of splenic follicular architecture in volasertib-treated mice in comparison to controls (**Figure 7C**), along with a partial restoration of normal hematopoietic islands in BM of volasertib-treated mice in comparison to vehicle-treated controls (**Figure 7D**). We confirmed these findings by carrying out flow cytometry which also demonstrated a complete clearance of human leukocytes (hCD45 vs mCD45.1) from the PB, BM and spleens of volasertib treated mice, in comparison to vehicle-treated controls (**Figure 7E-F**). Median spleen sizes and volumes were also significantly lower in volasertib-treated mice compared to vehicle-treated controls (**Figure 7G-H**). Consequently, these data support pharmacological inhibition of PLK1 as a potentially efficacious therapeutic strategy in pCMML.

## DISCUSSION

Chronic myelomonocytic leukemia is an aggressive hematological malignancy with dismal outcomes, with the pCMML subtype in particular having median OS of <2 years and high rates of AML transformation (Patnaik et al., 2014). Given limited therapeutic options for affected patients, we carried out this study to define the genetic and epigenetic landscape of pCMML and identify therapies that could modify disease biology. Phenotypically related to pCMML, juvenile myelomonocytic leukemia (JMML), a pediatric neoplasm is often initiated by germline or somatic RAS-activating mutations and is described as a *bona fide* RASopathy (Rauen, 2013; Stieglitz et al., 2015a). JMML outcome is associated with a dose-dependent effect for RAS pathway activation and the accumulation of additional recurrent mutations in genes involved in splicing, polycomb repressive complex 2 (PRC2) and gene transcription (Caye et al., 2015; Stieglitz et al., 2015a; Stieglitz et al., 2015b). Here, we show in pCMML that acquisition of RAS pathway mutations often occurs early, commonly on a background of epigenetic and splicing gene mutations and correlates with WHO-defined prognostic factors including leukocytosis, BM blasts and LT to sAML; further implying that severe forms of this disease often seen in the elderly, are also RASopathies that contrary to JMML, develop on a background of age-related clonal alterations (Mason et al., 2016). Analysis of mouse models recently suggested that loss of *Tet2*, which induces hypermethylation of *Spry2* encoding a negative regulator of the RAS signaling pathway, results in a fully penetrant CMML phenotype when combined with *Nras^G12D^* expression in mouse hematopoietic cells to synergistically enhance MAPK signaling. These findings suggest that pre-existing mutations in *TET2* may favor expansion of cells that acquire a RAS pathway mutation (Kunimoto et al., 2018). Of note, hematopoietic progenitor colony hypersensitivity to GM-CSF, an established hallmark of JMML, has also been depicted in CMML, especially in RAS mutant pCMML (Padron et al., 2013) and was also seen in progenitor cells obtained from the novel *Nras*^G12D^-Vav-Cre pCMML mouse model in this study.

WES studies have shown that, on average, CMML patients harbor between 10-15 somatic mutations per exon/coding region, similar to *de novo* AML and other myeloid malignancies (Ball et al., 2016; Merlevede et al., 2016). Of these 10-15 somatic mutations, a mean number of three affect recurrently mutated genes. These later genes (∼40 have been identified) encode mostly epigenetic regulators (*TET2*-60%, *ASXL1*-40%), proteins of pre-mRNA splicing complexes (*SRSF2*-50%) and members of signaling pathways (*NRAS* 30%; JAK2 10%) (Patnaik et al., 2014; Patnaik et al., 2016; Patnaik and Tefferi, 2016). Previous analyses of clonal architecture at the single cell level have shown that, in most cases, oncogenic mutations of the RAS pathway were secondary events (Itzykson et al., 2013b; Merlevede et al., 2016). Accordingly, we detected an increased incidence of RAS pathway mutations, especially those affecting *NRAS*, in LT compared to paired chronic phase CMML and in pCMML compared to dCMML. Interestingly, *NRAS* and *PTN11* are among the recurrently mutated genes that were shown to be associated with faster transformation of myelodysplastic syndromes into AML (Makishima et al., 2017). Of note, compared to chronic phase CMML, sAML demonstrated additional somatic mutations in only 44% of cases, indicating that LT can occur without additional driver mutations in the coding regions of DNA. Acquisition of *RAS* mutations correlated with WBC proliferation and blast cell accumulation, which are the two main prognostic factors recognized by the WHO in CMML, suggesting that *RAS* mutations form the landscape for epigenetic and evolutionary events leading to AML transformation.

We show that acquisition of RAS pathway mutations in CMML samples is associated with an increased expression of *PLK1* and *WEE1*. These genes encode mitotic checkpoint serine/threonine kinases that are suitable for therapeutic targeting. Polo-like kinases are involved in the regulation of centrosome maturation, bipolar spindle formation, anaphase-promoting complex, and chromosome segregation (Liu et al., 2016), whereas WEE1 influences cell cycle and cell size by inhibiting Cyclin-Dependent Kinase 1 (CDK1) (Matheson et al., 2016). We document a decrease in CMML progenitor cell growth when depleting *PLK1*and *WEE1*, supporting the suitability of targeting these check point kinases in RAS-mutated CMML. Another gene whose expression is increased in RAS pathway mutated CMML is *KMT2A (MLL1)*, which encodes a critical regulatory lysine methyltransferase. This later enzyme drives the monomethylation of H3K4 histone mark at the promoters of *PLK1* and *WEE1*. H3K4me1 is an activating histone configuration associated with increased gene transcription. While the entire KMT2 family (KMT2A-D, SET1A and SET1B) can result in methylation of H3K4, only KMT2A and KMT2B do so using MEN1 as a coactivator (Kerimoglu et al., 2017). We demonstrate KMT2A to be overexpressed in RAS-mutated pCMML, along with co-occupancy of MEN1 with KMT2A at the *PLK1* and *WEE1* promoters. While KMT2C and KMT2D can result in a MEN1 independent monomethylation of H3K4, we detected no changes in H3K4me1 enrichment at the promotor sites of *PLK1* and *WEE1* using siRNA against these lysine methyltransferases in comparison to siNT controls. Unexpectedly, while KMT2A can cause trimethylation of H3K4 (Kerimoglu et al., 2017), using ChIP-seq, both globally and in a sequence specific manner, we did not see any difference in H3K4me3 enrichment between RAS pathway mutant and wildtype CMML samples. In addition, Mi-503, a small molecule inhibitor of the KMT2A-MEN1 interaction induces a decrease in the expression of *PLK1*and *WEE1*, along with a decreased occupancy of H3K4me1 at these gene promoters. While translocations and PTD of *KMT2A* have long been implicated in AML pathogenesis, less than 1% of CMML patients demonstrate *KMT2A* translocations (Patnaik et al., 2018b), while there were no patients with PTD, suggesting for the first time that an unmutated but overexpressed *KMT2A* can also have an oncogenic role (Borkin et al., 2015).

*TET2, ASXL1* and *SRSF2* are the most common gene mutations encountered in CMML, with loss of function *ASXL1* mutations being prognostically detrimental (Patnaik et al., 2014; Patnaik et al., 2016; Patnaik and Tefferi, 2018). *ASXL1* mutations are thought to decrease the catalytic activity of the PRC2, thus resulting in depletion of H3K27me3, a repressive mark, depletion of which results in transcriptional upregulation (Abdel-Wahab et al., 2012; Abdel-Wahab and Levine, 2013). A second school of thought attributes detrimental effects of *ASXL1* mutations to an enhanced activity of the ASXL1-BAP1 complex, resulting in global erasure of H2AK119Ub, and H3K27me3 (Balasubramani et al., 2015). By using ChIP-seq in CMML samples, we demonstrate that H3K27me3 occupancy and recruitment at *PLK1* and *WEE1* promoters is not significantly different in RAS pathway mutated pCMML in comparison to RAS pathway wildtype dCMML, regardless of the *ASXL1* mutational status (sequence specific analysis restricted to *PLK1* and *WEE1*). *Asxl1* ^G643fs^ ^het^ knock in mice develop low risk MDS like features with very few animals developing MDS/MPN overlap syndromes like CMML and without AML transformation (Uni et al., 2019). Similarly, by carrying out DIP-seq, we demonstrate that there were no differences in 5-mc or 5-hmc levels at the promotors of *PLK1* and *WEE1* in RAS pathway mutant pCMML and RAS pathway wild-type dCMML patient samples, regardless of their *TET2* mutational status (sequence specific analysis restricted to *PLK1* and *WEE1*).

Both PLK1 (volasertib) and WEE1 inhibitors (AZD-1775) are currently in clinical trials for patients with myeloid neoplasms with reasonable safety data, prompting us to establish the therapeutic efficacy of these agents in pCMML models (Caldwell et al., 2014; Dohner et al., 2014; Janning and Fiedler, 2014; Ottmann et al., 2018). While a recent randomized phase III clinical trial, testing the efficacy of volasertib with low dose cytarabine (LDAC vs LDAC and placebo-NCT01721876) in AML patients ineligible for standard induction chemotherapy, did not meet its stated endpoints, this trial was not personalized to subsets of AML patients more likely to respond to therapy. In this study, the overall response rates for volasertib with LDAC vs LDAC with placebo were 25% and 16%, respectively (p=0.07). Efficacy data with regards to the use of the WEE1 inhibitor (AZD-1775) in combination with azacitidine in myeloid malignancies is currently not available. We show that targeting PLK1with volasertib, both *in vitro* and *in vivo*, results in effective killing of neoplastic cells, restoring normal hematopoiesis, with a differential sensitivity favoring *RAS* mutant pCMML cells. Given that these drugs have an established safety record in myeloid malignancies such as MDS and AML, we aim to translate these findings into an early phase clinical trial for pCMML.

In summary, we demonstrate that acquisition of oncogenic RAS pathway mutations defines a unique subgroup of CMML with poor outcomes. The detection of these mutations also offers a therapeutic window of opportunity for pCMML by the identification of the significance of KMT2A expression and downstream mitotic check point kinases PLK1 and WEE1, which can be targeted efficiently using existing small molecule inhibitors. These novel findings provide detailed insight into the biology of CMML and lay the foundation for the development of personalized clinical therapeutics for this otherwise devastating disease.

## ACKNOWLEDGEMENTS

Current publication is supported in part by grants from the “Gerstner Family Career Development Award”, The “Center for Individualized Medicine-Mayo Clinic”, and “The Henry J. Predolin Foundation for Research in Leukemia, Mayo Clinic, Rochester, MN, USA”.

This publication was also supported by CTSA Grant Number KL2 TR000136 from the National Center for Advancing Translational Science (NCATS). Its contents are solely the responsibility of the authors and do not necessarily represent the official views of the NIH.

S.I.N. was supported by Foundation ARC 2017, Foundation Gustave Roussy and Swiss Cancer League (grant KFC-3985-08-2016) grants. ES group is labelled “Equipe labellisée de la Ligue Nationale Contre le Cancer and supported by the French National Cancer Institute.

P.V. was supported by the Austrian National Science Fund (FWF) grants F4704-B20 and P30625-B28. A.A.M was supported by the Conquer Cancer Foundation of American Society of Clinical Oncology (ASCO) Young Investigator Award (YIA) and Grant. We would like to thank the University of Wisconsin Carbone Comprehensive Cancer Center (UWCCC) for use of its Shared Services (Flow Cytometry Laboratory, Genome Editing and Animal Models Shared Resource, and Experimental Pathology Laboratory) to complete this research. This work was supported by R01CA152108 and a Scholar Award from the Leukemia & Lymphoma Society to J.Z. This work was also supported in part by NIH/NCI P30 CA014520-UW-Comprehensive Cancer Center Support.

## AUTHOR CONTRIBUTIONS

R.M.C., D.V., T.L., D.M., E.J.T., L.L.A., G.C., I.H., A.B., A.M., S.N. and M.M.P. performed experiments, analyzed sequencing data and helped write the manuscript. A.V., X.Y., M.B., E.P. and J.J. helped perform mouse experiments with A.V., M.B. and E.P. conducting PDX experiments and X.Y. and J.J. developing the GEMM. B.A., K.R. and M.B. helped with DIP-seq. C.P., K.P., M.F.Z. and M.B. helped with ChIP-seq. M.H. helped with histopathology and flow cytometry. G.S., P.V., T.G., K.G. and M.M.P. helped write and edit the manuscript.

## METHODS

### Authorizations, patient cohorts, cell collection and sorting

This study was carried out at the Mayo Clinic in Rochester, Minnesota, after due approval from the Mayo Clinic Institutional Review Board (IRB-15-003786), and at Gustave Roussy Cancer Center in Villejuif, France, following the approval of the ethical committee Ile-de-France 1 (DC-2014-2091). Animal experiments were carried out at the Mayo Clinic after approval from the Mayo Clinic IACUC (protocol A00003488-18), the Moffitt Cancer Center in Florida (protocol) and the University of Wisconsin at Madison (protocol). Several patient cohorts were analyzed. In all cases, diagnosis was according to 2016 iteration of the WHO classification of myeloid malignancies (Arber et al, 2016). Cohort # 1 included 48 CMML patients whose paired MNC samples were collected in chronic phase and in LT to perform whole exome sequencing (WES). Cohort # 2 of 18 CMML patients was used to perform WES on samples serially collected along disease progression. This cohort had been described previously (Merlevede et al, 2016). Cohort # 3 was made of 576 CMML patients collected at Mayo Clinic; genetic abnormalities were screened in 350 of these patients using a 36-gene panel targeted next-generation sequencing (NGS) assay. Cohort # 4 was made of 592 CMML patient samples collected in France (n=417) and in Austria (n=175) with molecular information obtained by NGS using a 38-gene panel targeted NGS assay. PB and BM samples were collected in EDTA tubes after due informed consent. MNCs were collected on Ficoll-Hypaque whereas CD14- and CD3-positive cells were sorted using magnetic beads and the AutoMacs system (Miltenyi Biotech, Bergish Gladbach, Germany). In 5 cases, buccal mucosa cells were used as germline control cells.

### Whole Exome Sequencing (WES)

One μg of genomic DNA was sheared with the Covaris S2 system (LGC Genomics). Exome capture and sequencing library preparation were performed using Agilent SureSelect Human All Exon V5 (Agilent Genomics, Santa Clara, CA), following standard protocols. The final libraries were indexed, pooled and paired-ends (2 × 100 bp) sequenced on Illumina HiSeq platforms (San Diego, CA). Germ line DNA was obtained from CD3-positive cells for 5 patients of the CMML LT dataset and 17 patients with serial WES analyses (Merlevede et al., 2016). The mean coverage in the targeted regions was 100-fold. Sequencing data are deposited at the European Genome–Phenome Archive (EGA), hosted by the European Bioinformatics Institute (EBI). Paired-end reads in the form of FASTQ files were aligned to the GRCh37 version of the human genome with BWA or Novalign to generate BAM files. Variant calling analysis was performed using Agilent SureCall software. Genomic variant information including evidence of and somatic status in cancer, significant protein domains, and known pathogenicity was further annotated using public databases: the Catalogue of Somatic Mutations in Cancer - COSMIC (https://cancer.sanger.ac.uk/cosmic), ClinVar (https://www.ncbi.nlm.nih.gov/clinvar) and UniProt (https://www.uniprot.org). Allele frequency in the normal population was calculated using the Exome Aggregation Consortium ExAC (http://exac.broadinstitute.org). Alternative alleles with less than five reads and/or a frequency less than 5% were excluded. All variants passing the sequencing quality-control filters and within a gene coding region, regardless of database population frequency, underwent characterization. Since CD3-positive cells are known to be contaminated with tumor cells we performed MuTect2 in the ‘tumor-only’ mode (Stieglitz et al., 2015a). For all identified mutations and indels variant allele frequencies (VAF) in the corresponding CD3-positive DNA was retrieved from BAM files. All the variants with p values from Fisher exact test below 10^-4^ were considered as somatic. Regions of somatic copy number alterations (SCNA) were screened manually. A panel of normal samples (PON) was constructed from 50 germline samples and was used to reduce the rate of false positives. Additionally, only stop-gain, splice-site and probably damaging nonsynonymous mutations were considered to be putative drivers in tumor suppressor genes. SCNA analysis was performed with Facets statistical and computational analysis pipeline in ‘tumor-normal’ mode where possible, or in ‘tumor only’ mode. cnLOH was called based on the germline heterozygous SNPs VAFs in the tumor (Shen and Seshan, 2016). Tumor evolution was visualized with ClonEvol software (https://ieeexplore.ieee.org/document/6650525).

### Targeted Next Generation Sequencing (NGS)

Analysis of the Mayo cohort (#3) was done using a panel in which the regions of 36 genes was selected for custom target capture using Agilent SureSelect Target Enrichment Kit. Briefly, libraries derived from each DNA sample were prepared using NEB Ultra II (New England Biolabs, Ipswitch, MA) and individually bar coded by dual indexing. Sequencing was performed on a HiSeq 4000 (San Diego, CA) with 150-bp paired-end reads and consisted of 48 pooled libraries per lane. Median sequence read depth was ∼400×. The custom panel of target regions covered all coding regions and consensus splice sites from the following 36 genes: *ASXL1, CALR, CBL, CEBPA, DNMT3A, EZH2, FLT3, IDH1, IDH2, IKZF1, JAK2, KRAS, MPL, NPM1, NRAS, PHF6, PTPN11, RUNX1, SETBP1, SF3B1, SH2B3, SRSF2, TET2, TP53, U2AF1,* and *ZRSR2.* Paired end reads (FASTQ files) were processed and analyzed using same method as WES and as previously published by our group (Patnaik et al., 2017; Patnaik et al., 2014; Patnaik et al., 2016). The French part of cohort (# 4) was analysed using an Ion AmpliSeq™ Custom Panel Primer Pools (10 ng of gDNA per primer pool) to perform multiplex PCR targeting all coding regions or selected regions of the following 36 genes (92 kb): *ASXL1* (exons 11-12), *BCOR* (2-14), *BRAF* (15), *CBL* (4-5, 7-9, 11-14), *CEBPA* (1), *CSF3R* (3-18), *DNMT3A* (2-23), *ETV6* (1-8), *EZH2* (2-20), *FLT3* (15-20), *GATA2* (4-6), *IDH1* (4, 6), *IDH2* (4), *JAK2* (12, 14), *KDM6A* (full), *KIT* (2, 8-11, 13-14, 16-17), *KRAS* (2-4), *MPL* (10), *MYD88* (5), *NRAS* (2-4), *OGT* (1-22), *PHF6* (1, 4, 6-7, 9), *PTPN11* (1-15), *RAD21* (2-14), *RIT1* (4-5), *RUNX1* (2-9), *SETBP1* (4-5), *SF3B1* (full), *SRSF2* (1-2), *STAG1* (*2-34*), *STAG2* (3-35), *TET2* (3-11), *TP53* (2-11), *U2AF1* (2, 6), *WT1* (1, 7) and *ZRSR2* (full). PCR amplicons after Fupa digestion were end-repaired, extended with an ‘A’ base on the 3′end, ligated with indexed paired-end adaptors (NEXTflex, Bioo Scientific) using the Bravo Platform (Agilent), amplified by PCR for 6 cycles and purified with AMPure XP beads (Beckman Coulter). Sequencing (2x250 bp reads) was performed on a MiSeq flow cell (Illumina, San Diego, CA) using the onboard cluster method.

Sequencing reads were mapped to the GRCh37 version of reference genome with the Burrows-Wheeler Aligner (BWA) (PMID: 19451168) and then processed with the Genome Analysis Toolkit (GATK; version 4.1.2.0) (PMID: 20644199) following best practices for exomes and specific indications for smaller targeted regions. Sequencing quality and target enrichment were verified with Picard tools metrics. Germinal DNA was obtained from CD3-positive cells for 5 patients from the CMML/sAML dataset and for 17 patients from CMML with serial WES analyses. Since CD3-positive cells cells are known to be contaminated with tumor cells (PMID: 26457647; PMID: 26908133) we performed MuTect2 in the ‘tumor-only’ mode. For all identified mutations and indels Variant Allele Frequencies in the corresponding CD3+ DNA was retrieved from bam files. All the variants with p values from Fisher exact test below 10^-4^ were considered as somatic. Regions of Somatic copy Number Alterations SCNA were screened manually. A panel of normal samples (PON) was constructed from 50 germline samples ad was used to reduce the rate of false positives. We filtered out all polymorphisms from ExAC database (http://exac.broadinstitute.org) with allelic frequency above 0.001. Additionally, variants were considered putative drivers if reported in the COSMIC v89 database. Only stop-gain, frameshift indels, splice-site and probably damaging nonsynonymous mutations were considered to be putative drivers in tumor suppressor genes. SCNA analysis was performed with Facets statistical and computational analysis pipeline in ‘tumor-normal’ mode where possible, or in ‘tumor only’ mode. cnLOH was called based on the germline heterozygous SNPs VAFs in the tumor (PMID: 27270079).

All the variant processing was conducted in python with pandas library. Fisher exact test was used for contingency tables. Mann-Whitney test was used for two groups of continuous variables. Pearson-Spearman test was used for pairwise correlations of continuous variables. Log-rank test was used for survival analyses. Multivariate analysis was executed using logistic regression model, LASSO with selected features (Regression Shrinkage and Selection via the Lasso. Robert Tibshirani) and applied to measure enrichment / depletion between two subgroups. Tumor evolution was visualized with ClonEvol software (https://ieeexplore.ieee.org/document/6650525).

### Hematopoietic progenitor colony forming assay

PB and/or BM samples were first treated with ammonium chloride to deplete red blood cells, washed in RPMI, then plated at a final concentration of 5x10^4^-2x10^5^ cells/mL into methylcellulose formulated with recombinant cytokines to support the optimal growth of erythroid progenitor cells (BFU-E and CFU-E), granulocyte-macrophage progenitor cells (CFU-GM, CFU-G and CFU-M), and multipotent granulocyte, erythroid, macrophage and megakaryocyte progenitor cells (CFU-GEMM). Plates were incubated at 37°C and colonies were enumerated on day 10-14 as described (Pardanani et al., 2007). Individual colonies were isolated, placed into a 40mM NaOH buffer, and heated to 95°C for 15 min. Up to 5uL of product was submitted for polymerase chain reaction (PCR) using standard conditions. Cycling conditions consisted of: an initial denaturation at 94°C for 2 min; 40 cycles of denaturation at 94°C for 30 sec, annealing at 56°C for 30 sec, and extension at 72°C for 1 min, and final extension at 72°C for 3 minutes. PCR products were visualized by 1.3% agarose gel, purified, and the final product was subjected to bidirectional sequence analysis (GeneWiz, South Plainfield, NJ).

### RNA Sequencing

Library preparation for the RNA samples was done using Illumina TruSeq Stranded Total RNA and sequencing was done using Illumina HiSeq 4000, 100 cycles x 2 paired-end reads. RNA-seq sequencing reads were processed through the MAPRSeq v2.0. bionformatics workflow as described in(Kalari et al., 2014). Briefly reads were mapped using TopHat version 2.06 against the reference GRCh37 without alternative haplotypes using the transcript models from UCSD (March 2012) available from Illumina iGenomes Project. Gene counts were calculated with featureCounts (Liao et al., 2014), differential expression was performed using edgeR and clustering was performed using heatmap.2(Robinson et al., 2010). RNA samples were prepared and RNA quality was determined using an Agilent Bioanalyzer RNA Nanochip or Caliper RNA assay and arrayed into a 96-well plate. The paired-end sequencing libraries were prepared following BC Cancer Genome Sciences Centre’s strand-specific, plate-based library construction protocol on a Microlab NIMBUS robot (Hamilton Robotics, USA). Libraries were sequenced on an Illumina HiSeq2500 platform to a read depth of approximately 50 million reads per sample. Reads were aligned to the human genome (GRCh38/hg38) using the HISAT2 aligner (Kim et al., 2015). Read counts were generated using featureCounts with reference to Homo_sapiens.GRCh38.90.gtf annotation file from Ensembl (Liao et al., 2014; Zerbino et al., 2018). Patient’s samples were grouped based on the previously identified histological classifications and differential expression analyses were performed using the R Bioconductor package DESeq2 (Love et al., 2014). Pathway data was derived from ConsensusPathDB (Kamburov et al., 2009). Heatmaps were derived using the R package pheatmap (pheatmap: CRAN.R-project.org/package=pheatmap) and principle component analyses were derived using the R package rgl (rgl: CRAN.R-project.org/package=rgl)

### RNA PCR

For the RNAseq validation we analyzed a subset of target genes by Quantitative Reverse Transcription Polymerase Chain Reaction (RT-qPCR). Total left-over RNA from the RNAseq was used to synthetize cDNA with the High-Capacity cDNA Reverse Transcription Kit (Applied Biosystems, Carlsbad, CA). A dilution 1/3 of the total cDNA was amplified by real-time PCR. Samples were prepared with PerfeCTa SYBR Green FastMix (Quanta BioSciences Inc., Gaithersburg, MD) and the following primer sets:

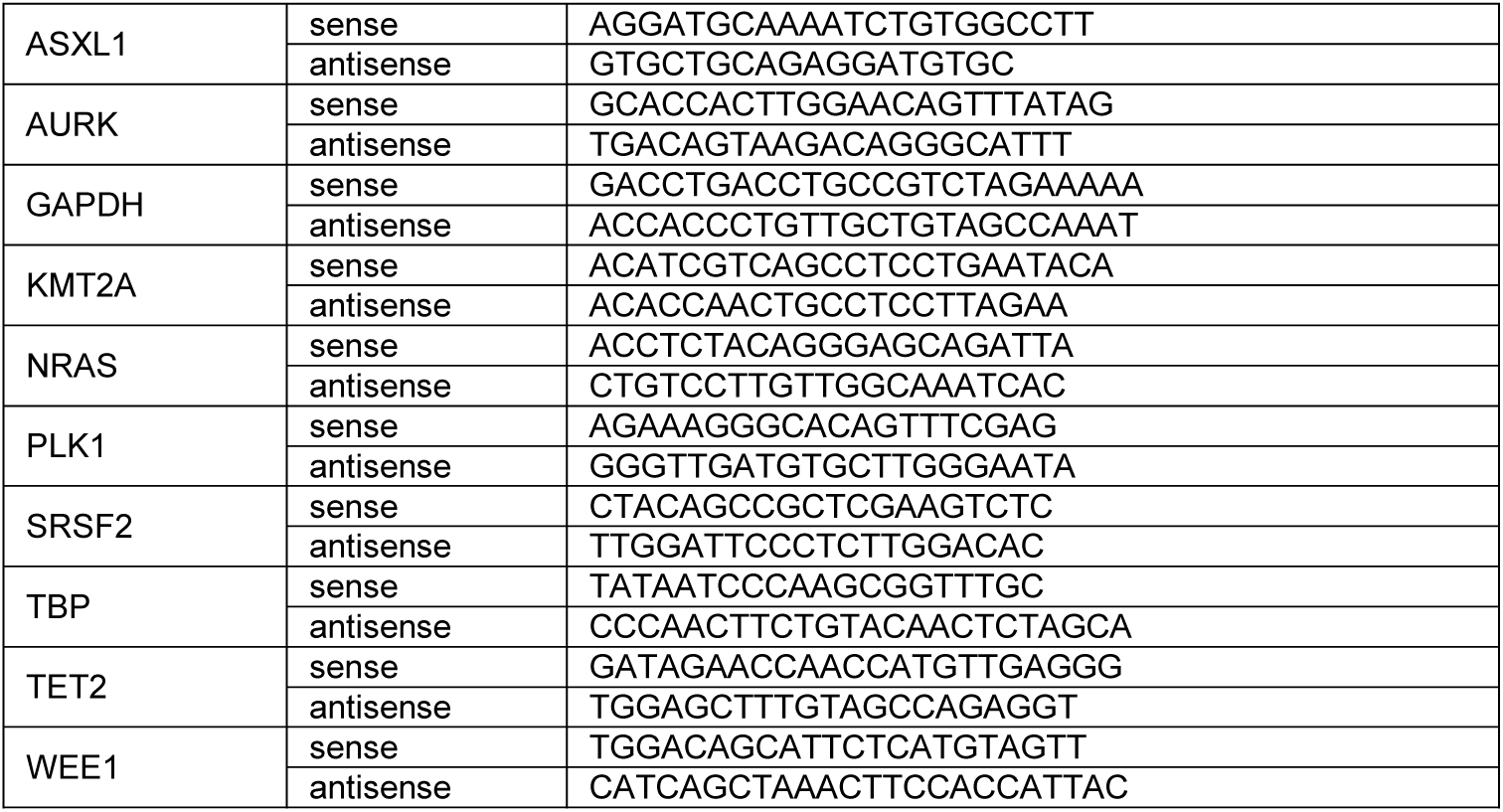

### Chromatin immunoprecipitation and sequencing (ChIP-seq)

Approximately 100,000 cells from each CMML patient sample were used for input for native chromatin immunoprecipitation (N-ChIP). Cells were lysed on ice for 20 minutes in lysis buffer containing 0.1% Triton X-100, 0.1% deoxycholate and protease inhibitor. Extracted chromatin was digested with 90U of MNase enzyme (New England Biolabs) for 6 minutes at 25°C. The reaction was quenched with 250 µM of EDTA post-digestion. A mix of 1% Triton X-100 and 1% deoxycholate was added to digested samples and incubated on ice for 20 min. Digested chromatin was pooled and pre-cleared in IP buffer (20 mM Tris-HCl [pH7.5], 2 mM EDTA, 150 mM NaCl, 0.1% Triton X-100, 0.1% deoxycholate) plus protease inhibitors with pre-washed Protein A/G Dynabeads (Thermo Fisher Scientific; Waltham, MA, USA) at 4°C for 1.5 hours. Supernatants were removed from the beads and transferred to a 96-well plate containing the antibody-bead complex. Following an overnight 4°C incubation, samples were washed twice with Low Salt Buffer (20 mM Tris-HCl [pH 8.0], 0.1% SDS, 1% Triton X-100, 2 mM EDTA, 150 mM NaCl) and twice with High Salt Buffer (20 mM Tris-HCl [pH 8.0], 0.1% SDS, 1% Triton X-100, 2 mM EDTA, 500 mM NaCl). DNA-antibody complexes were eluted in Elution Buffer (100 mM NaHCO_3_, 1% SDS), incubated at 65°C for 90 minutes. Protein digestion was performed on the eluted DNA samples at 50°C for 30 minutes (Qiagen Protease mix). ChIP DNA was purified using Sera-Mag beads (Fisher Scientific) with 30% PEG before library construction. Library construction was prepared by following a modified paired-end library protocol (Illumina Inc., USA) and sequenced on an Illumina HiSeq2500 sequencer to a depth of ∼ 25 million (H3K4me1 and H3K4me3) or ∼50 million reads (H3K27me3 and Input) per sample. Reads were aligned to the human genome (GRCh38/hg38) using bowtie2 (Langmead and Salzberg, 2012). Aligned reads were sorted and indexed using samtools (Li et al., 2009) and peaks were called using macs2 (Zhang et al., 2008). Regions of differential enrichment were derived using the R Bioconductor package DiffBind (*DiffBind: differential binding analysis of ChIP-Seq peak data*. http://bioconductor.org/packages/release/bioc/vignettes/DiffBind/inst/doc/DiffBind.pdf) and areas of interest were annotated using Homer (Ramirez et al., 2016). Track files were generated using deeptools and visualized on the UCSC Genome Browser (Kent et al., 2002).

### Chromatin immunoprecipitation-qPCR

ChIP was performed as described previously (Comba et al., 2016). Briefly, PB MNC from CMML patients (20 x 10^6^) were crosslinked with 1% formaldehyde directly into the medium for 10 min at room temperature. The cells were then washed with PBS, collected by centrifugation at 800 x *g* for 5 minutes at 4 °C, re-suspended in Cell Lysis Buffer (20mM Tris [pH 8];15mM PIPES; 85mM KCl; 0.5% NP-40), and incubated on ice for 15 minutes. Resulting pellets were then re-suspended in Nuclear Lysis Buffer (50mM Tris [pH 8.1]; 10mM EDTA; 1% SDS) and chromatin was sheared to an average fragment size of 200-600 bp using a water bath sonicator (Bioruptor, Diagenode, Denville, NJ). Input chromatin was retained from each individual sample prior to addition of antibody for qPCR analysis. Chromatin was incubated overnight at 4°C with magnetic beads (Dynabeads Protein G, Invitrogen) and the following antibodies (4 µg each): KMT2A (Abcam); MEN1(Abcam); H3K4me3 (EMD Millipore); H3K4me1 (Abcam); normal rabbit IgG (EMD Millipore) and normal mouse IgG (EMD Millipore). Following immunoprecipitation, crosslinks were removed, and immunoprecipitated DNA was purified using spin columns and subsequently amplified by qPCR. Quantitative PCR was performed using primer sets to amplify different regions of the human PLK1 and WEE1 promoters (for primers sequences and position on the promoter see Table below). Quantitative SYBR PCR was performed in triplicate for each sample or input using the C1000 Thermal Cycler (BioRad).

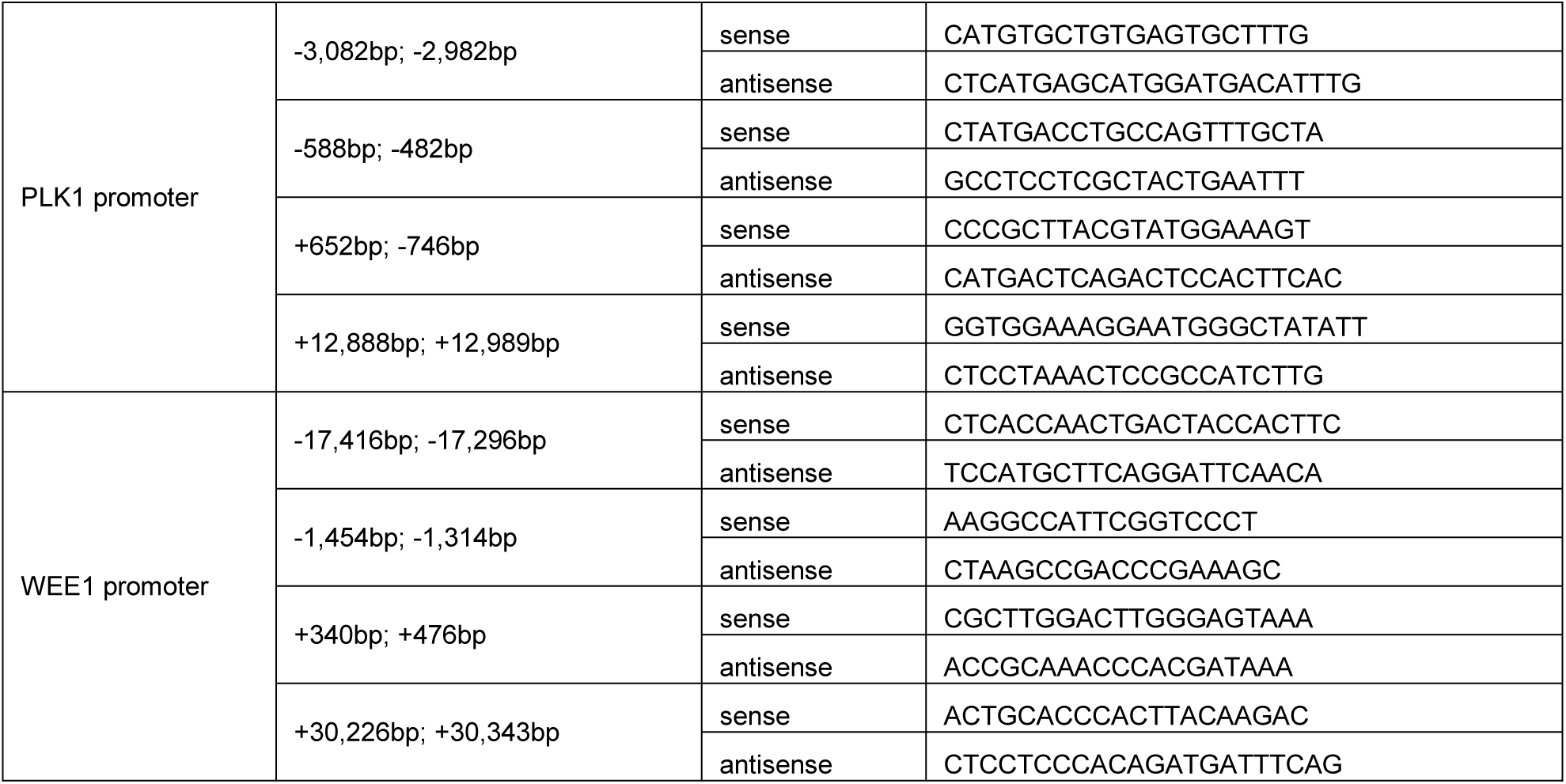

### DNA immunoprecipitation (DIP) -seq analysis of 5mC, and 5hmC DNA Marks

Genomic DNA was isolated from patient samples and submitted to the Mayo Clinic Epigenomics Development Laboratory (EDL) for DNA immunoprecipitation and library preparation (Swift bioscience ACCEL-NGS® 1S Plus DNA Library Kit). Antibodies used for this study included: 5mC-33D3 monoclonal antibody (Diagenode C15200081), 5hmC antibody (developed in-house by EDL), and bridging antibody (Active Motif 53017). HiSeq4000 paired-end sequencing was performed at the Medical Genome Facility at Mayo Clinic Rochester. Raw reads were mapped to the hg38 human reference genome using Burrows-Wheeler Aligner (BWA)(Li and Durbin, 2009). Averaged dysplastic vs. proliferative sample groups were prepared using Samtools merge of the corresponding aligned bam files. Averaged patient sample groups were as follows: Proliferative: SpecID 4, 11, 15, 18, 20, 21, 24, 26, & 29 and Dysplastic: SpecID 2, 5, 6, 8, 16, 22, 25, 27, & 28. Input corrected bigwig files were constructed using Deeptools bamcoverage and bigwigCompare (input subtraction correction) with UCSC hg38 gene coordinates (Ramirez et al., 2016). Tag density plots were constructed using Deeptools computeMatrix and PlotProfile. PB MNC H3K27ac ChIP-seq data was downloaded from GEO (GSM1127137) (Bernstein et al., 2010).

### Electroporation technique for siRNA experiments

Patient-derived PB and/or BM MNC were first treated with ammonium chloride to deplete mature, non-nucleated red blood cells, washed in RPMI, and then plated into methylcellulose (see Hematopoietic progenitor colony forming assays for further details) to allow for progenitor expansion. Cells were liberated from methylcellulose by solubilizing them with PBS warmed to 37 degrees C. Cells were washed with RPMI before being re-suspended in Lonza electroporation solution from Kit C (Lonza, Basel, Switzerland) and electroporated using program W-001 using the Amaxa nucleofector (Lonza, Basel, Switzerland). For every one million cells transfected, 2µg of plasmid constructs or 60pg of siRNA was used. Electroporation efficiency was monitored by transfecting an eGFP construct as a control. After electroporation, cells were maintained in RPMI supplemented with StemSpan (StemCell Technologies #02691) for 72 hours before experiments were conducted.

### Recombinant adenoviral transduction procedure

We utilized eGFP Adenovirus (#1060) and *NRAS* viral oncogene homolog adenovirus (#1565), both from Vector Biolabs (Malvern, PA) for the transduction procedure. Briefly, we aliquoted 10 million PBMC cells from CMML patients into 1 mL of RPMI and added either control (GFP) or *NRAS* expressing adenovirus at an MOI of 100. We spun the cells for 30 minutes at 1200 RPM and let the cells sit an additional 30 minutes at room temperature. We then collected the cells and plated them at the same concentration (using RPMI) into cytokine enriched methylcellulose media for colony formation (MethoCult^TM^GF H4434, StemCell Technologies, Vancouver, CA). After 7 days at 37°C, we enumerated and imaged the progenitor colony growth.

### *In vitro* drug studies on progenitor colonies with PLK1, WEE1 and KMT2A-MENIN1 inhibitors

Patient-derived PB and/or BM MNC were plated at 4 x 10^4^ cells per well in 96-well plates in 200 µL of RPMI supplemented with 10% fetal bovine serum and 10X StemSpan (StemCell Technologies #02691) (Pardanani et al., 2007). Media was also supplemented with either DMSO as a vehicle control of one the various inhibitors tested. AZD-1775 (WEE1 inhibitor) was tested at concentrations 10 nM, 50 nM, 100 nM, 250 nM and 500 nM. Volasertib (PLK1 inhibitor) was tested at concentrations 10 nM, 100 nM, 250 nM, 500 nM and 1000 nM. On day four after plating the cells, they were centrifuged to aspirate and replace media. Drug assays were completed on day seven. At this time, cells in each well were counted using a hemocytometer. Subsequently, an MTS assay was performed following the manufacturer’s protocol.

### Western blot

PVDF membranes were blocked in Tris-buffered saline and 0.3% Tween-20 (TBST) with 5% skim milk overnight at 4 °C. Primary antibodies used in western blotting were prepared in a solution of TBST with 2% skim milk at a working concentration of 1:1000 and included antibodies against PLK1 (4535), WEE1 (13084) and KMT2A (14197), and phospho-ERK1/2 (Thr202/Tyr204) all from Cell Signaling Technology. Antibodies were also used against vinculin (Bethyl, A302-535A) and total ERK1/2 (R&D, MAB1576). In order to assess volasertib-mediated mitotic kinase inhibition we used antibodies against histone 3 (4499), CDK1 (28439) and phospho-CDK1 Y15 (4539) all from Cell Signaling and phospho-histone 3 S10 (Millipore, 06-570). HRP-conjugated goat anti-rabbit IgG (Millipore, 12-348) and goat anti-mouse IgG (Millipore, 12-349) secondary antibodies were used at a concentration of 1:5000 and detected using SuperSignal West Dura Extended Duration Substrate (Thermo Scientific, 34076).

### Genetically engineered mouse models

The *NRAS*^G12D+^ genetically engineered mouse model used for this manuscript was developed in collaboration with the Zhang laboratory at the University of Wisconsin in Madison. All mouse lines were maintained in a pure C57BL/6 genetic background (>N10). Mice bearing a conditional oncogenic *Nras* allele (*Nras^Lox-stop-Lox^ ^(LSL)^ ^G12D/+^*) were crossed to *Vav-Cre* (Damnernsawad et al., 2016) mice to generate the experimental mice, including *Nras ^LSL^ ^G12D/+^; Vav-Cre*, and *Vav-Cre* mice. These mice were genotyped as previously described (Abdel-Wahab et al., 2013; Wang et al., 2011). All animal experiments were conducted in accordance with the *Guide for the Care and Use of Laboratory Animals* and approved by an Animal Care and Use Committee. For lineage analysis of PB and BM, flow cytometric analyses were performed as described (You et al., 2018). Stained cells were analyzed on a LSR II flow cytometer (BD Biosciences). Antibodies specific for surface antigens Mac-1 (M1/70) and Gr-1 (RB6-8C5) were purchased from eBioscience. Surface proteins were detected with FITC-conjugated antibodies (eBioscience unless specified) against B220 (RA3-6B2), Gr-1 (RB6-8C5), CD3 (17A2, Biolegend), CD4 (GK1.5), CD8 (53-6.7), and TER119 (TER-119), and PE-conjugated anti-CD117/c-Kit (2B8) antibody. To analyze phospho-ERK1/2, total BM cells were deprived of serum and cytokines and then stimulated with cytokines as previously described (You et al., 2018). Phosphorylated ERK1/2 (p-ERK1/2) was analyzed in defined Lin^-/low^ c-Kit^+^ cells as described. p-ERK1/2 was detected by a primary antibody against p-ERK (Thr202/Tyr204; Cell signaling Technology) followed by APC-conjugated donkey anti-rabbit F(ab’)2 fragment (Jackson ImmunoResearch, AB-2340601). Mouse tissues were fixed in 10% neutral buffered formalin (Sigma-Aldrich) and further processed at the Experimental Pathology Laboratory at the University of Wisconsin-Madison and at the Mayo Clinic.

### Patient derived xenograft (PDX) models

The NOD.Cg-*Prkdc^scid^ Il2rg^tm1Wjl^*/SzJ-SGM3 (NSGS) mice were used for the PDX experiments (Yoshimi et al., 2017). The mice were 6 to 8 weeks old at the start of the experiments with a weight of 20–30 grams. Mice were irradiated at 2.5 Gy with a cesium source irradiator and xenografted 24 hours after irradiation, with 1-5 x 10(6) BM MNCs from *NRAS* mutant CMML patients. Volasertib was dosed at 20 mg/kg body weight intraperitoneally once a week for 2 weeks (Gjertsen and Schoffski, 2015; Liu et al., 2016; Rudolph et al., 2015). Mice were monitored daily for signs of disease and euthanized by carbon dioxide at moribund appearance, or 2 weeks after the last treatment. Spleen, BM and PB cells were collected at the time of necropsy and processed for flow cytometry. Cells from blood, BM and spleen were stained with the following antibodies: murine CD45.1 BUV737 (BD Biosciences-564574), human CD45 BV605 (BD Biosciences-564048), human CD3 APC (BD Biosciences-561810) and human CD33 PE (BD Biosciences-555450). All blood, BM and spleen samples were analyzed on a LSRII flow cytometer (BD Biosciences). Femur, spleen, lung and liver samples were fixed overnight in 10% neutral buffered formalin, dehydrated, paraffin embedded and 5 micron sections were prepared. After hematoxylin-eosin staining was performed on all of the samples, sections were examined by a pathologist. Complete blood counts were also analyzed on all PDX mice at endpoint (IDEXX Veterinary Diagnostics). Immunohistochemistry staining was performed at the Pathology Research Core (Mayo Clinic, Rochester, MN) using the Leica Bond RX stainer (Leica). Slides for CD3 stain were retrieved for 30 minutes using Epitope Retrieval 2 (EDTA; Leica) and incubated in protein block (Rodent Block M, Biocare) for 30 minutes. The CD3 primary antibody (Clone F7.2.38; Dako) was diluted to 1:150 in Bond Antibody Diluent (Leica). Slides for CD14 and CD163 stains were retrieved for 20 minutes using Epitope Retrieval 2 (EDTA; Leica) incubated in protein block (Rodent Block M, Biocare) for 30 minutes. The CD14 primary antibody (Rabbit Polyclonal – HPA001887; Sigma) was diluted to 1:300 in Bond Antibody Diluent (Leica). The CD163 primary antibody (Clone 10D6, Leica) was diluted to 1:400 Background Reducing Diluent (Dako). All primary antibodies were incubated for 15 minutes. The detection system used was Polymer Refine Detection System (Leica). This system includes the hydrogen peroxidase block, post primary and polymer reagent, DAB, and Hematoxylin. Immunostaining visualization was achieved by incubating slides 10 minutes in DAB and DAB buffer (1:19 mixture) from the Bond Polymer Refine Detection System. To this point, slides were rinsed between steps with 1X Bond Wash Buffer (Leica). Slides were counterstained for five minutes using Schmidt hematoxylin and molecular biology grade water (1:1 mixture), followed by several rinses in 1X Bond wash buffer and distilled water, this is not the hematoxylin provided with the Refine kit. Once the immunochemistry process was completed, slides were removed from the stainer and rinsed in tap water for five minutes. Slides were dehydrated in increasing concentrations of ethyl alcohol and cleared in 3 changes of xylene prior to permanent cover slipping in xylene-based medium. Mice assessments included impact of therapy on the neoplastic clone (assessed by flow cytometry), impact on the percentage of circulating blasts and monocytes, impact on liver and spleen sizes and weights, impact on BM changes such as hypercellularity and dysplasia. We did also assess potential adverse events, including additional cytopenias, cutaneous and gastro intestinal complications. Volasertib dosing at 40 mg/kg body weight intraperitoneally did result in significant gastrointestinal toxicity (necrotic bowel loops) along with death of the treated mice.

### Statistical analysis

Distribution of continuous variables was statistically compared using Mann-Whitney or Kruskal-Wallis tests, while nominal or categorical variables were compared using the Chi-Square or Fischer’s exact test. Time to event analyses used the method of Kaplan-Meier for univariate comparisons using the log-rank test. OS was calculated from the date of diagnosis to date of death or last follow-up, while AML-free survival (LFS) was calculated from date of diagnosis to date of AML transformation or death.

