## Supplementary Information for "*RAS* mutations drive proliferative chronic myelomonocytic leukemia via activation of a novel KMT2A-PLK1 axis"

### Supplementary Figure 1

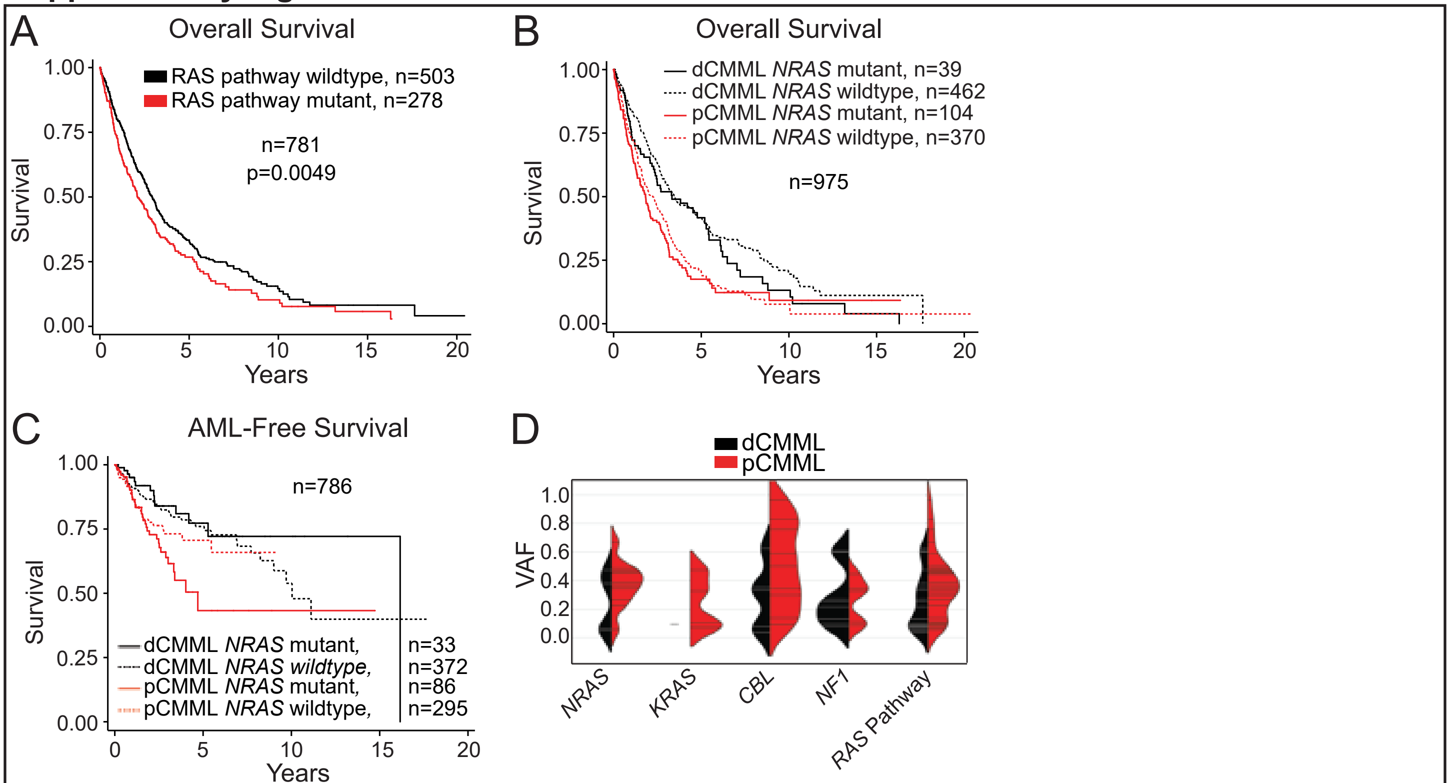

### Supplementary Figure 2

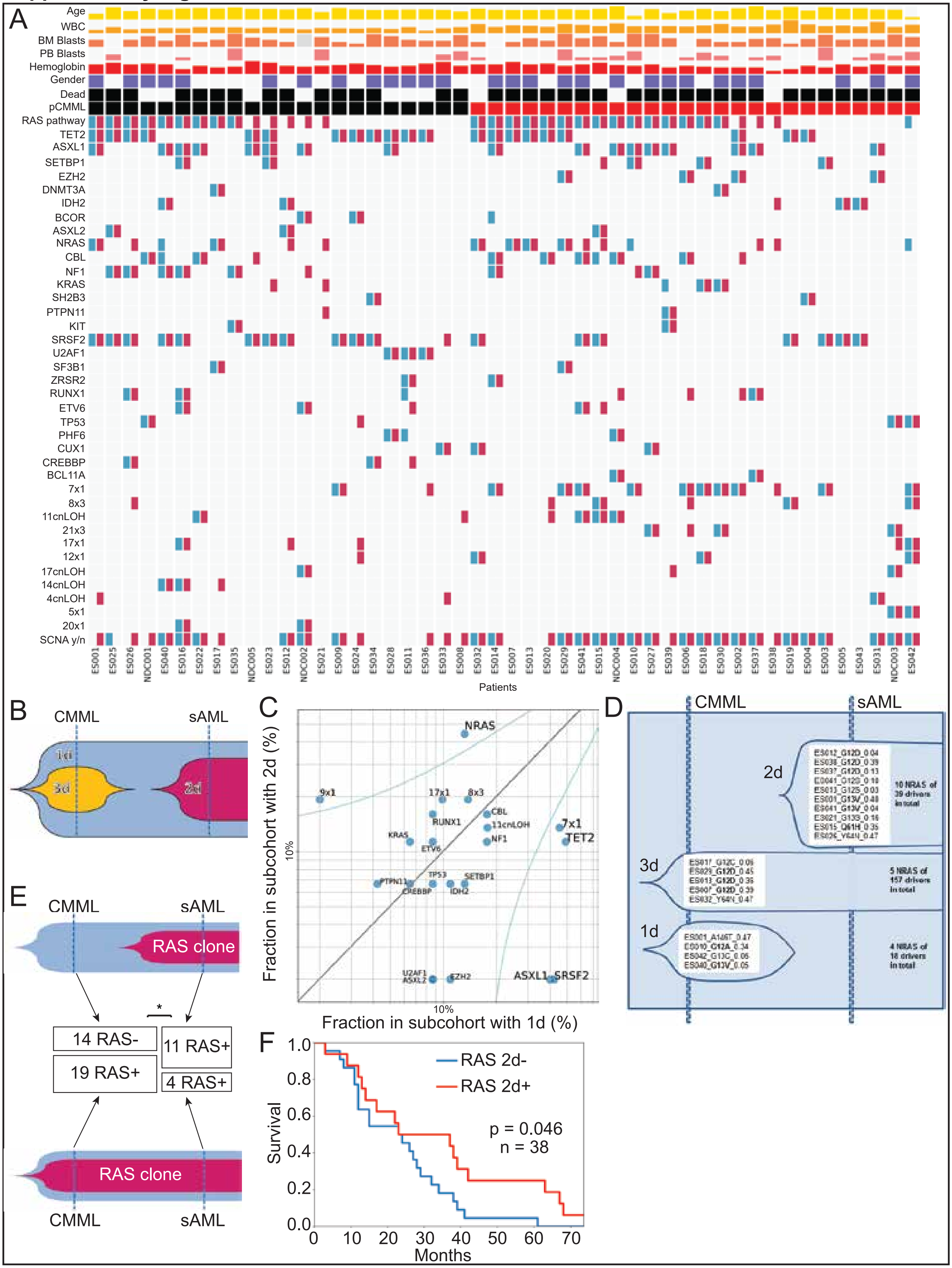

**Supplementary Figure 4**

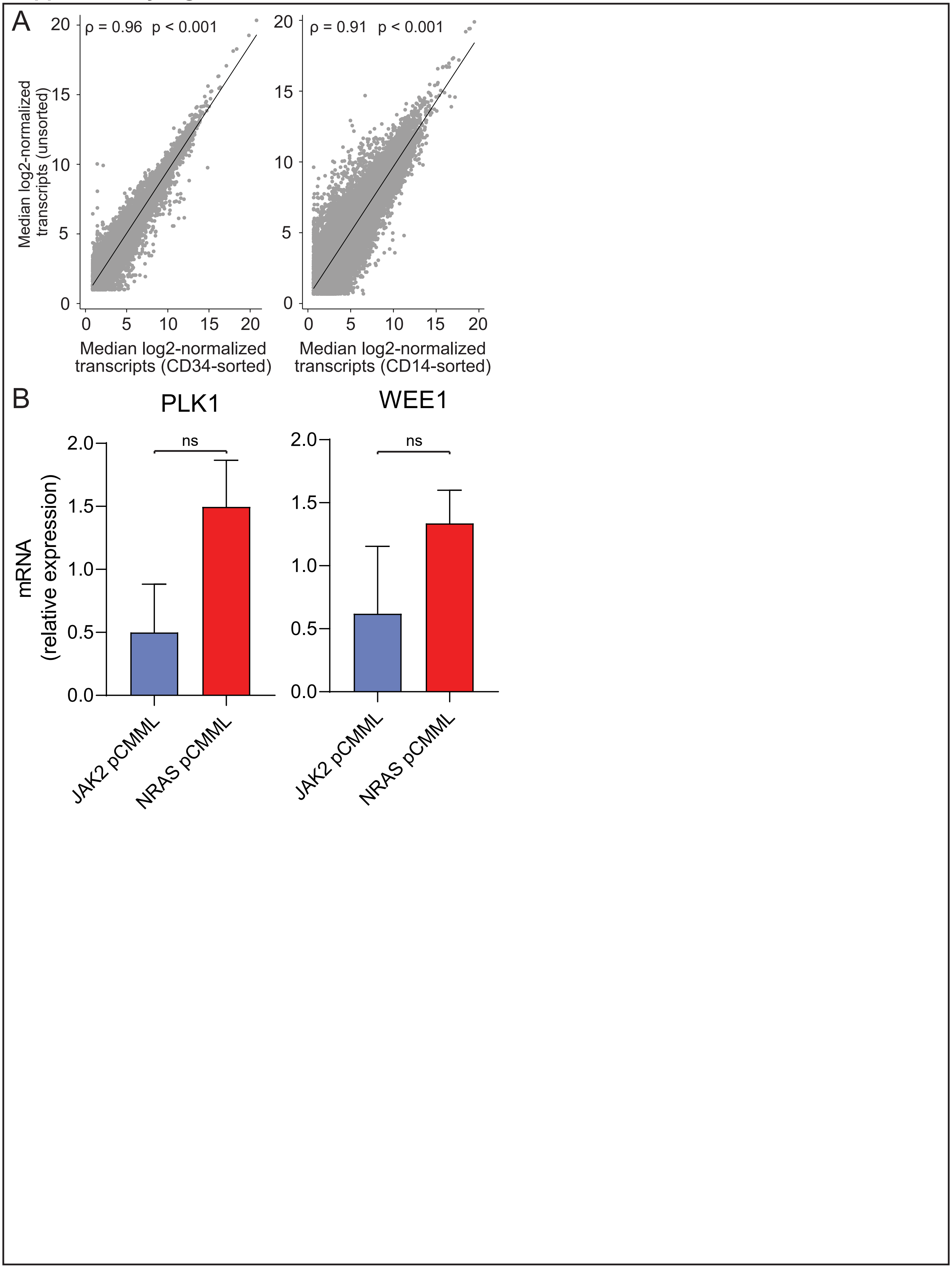

### Supplementary Figure 5

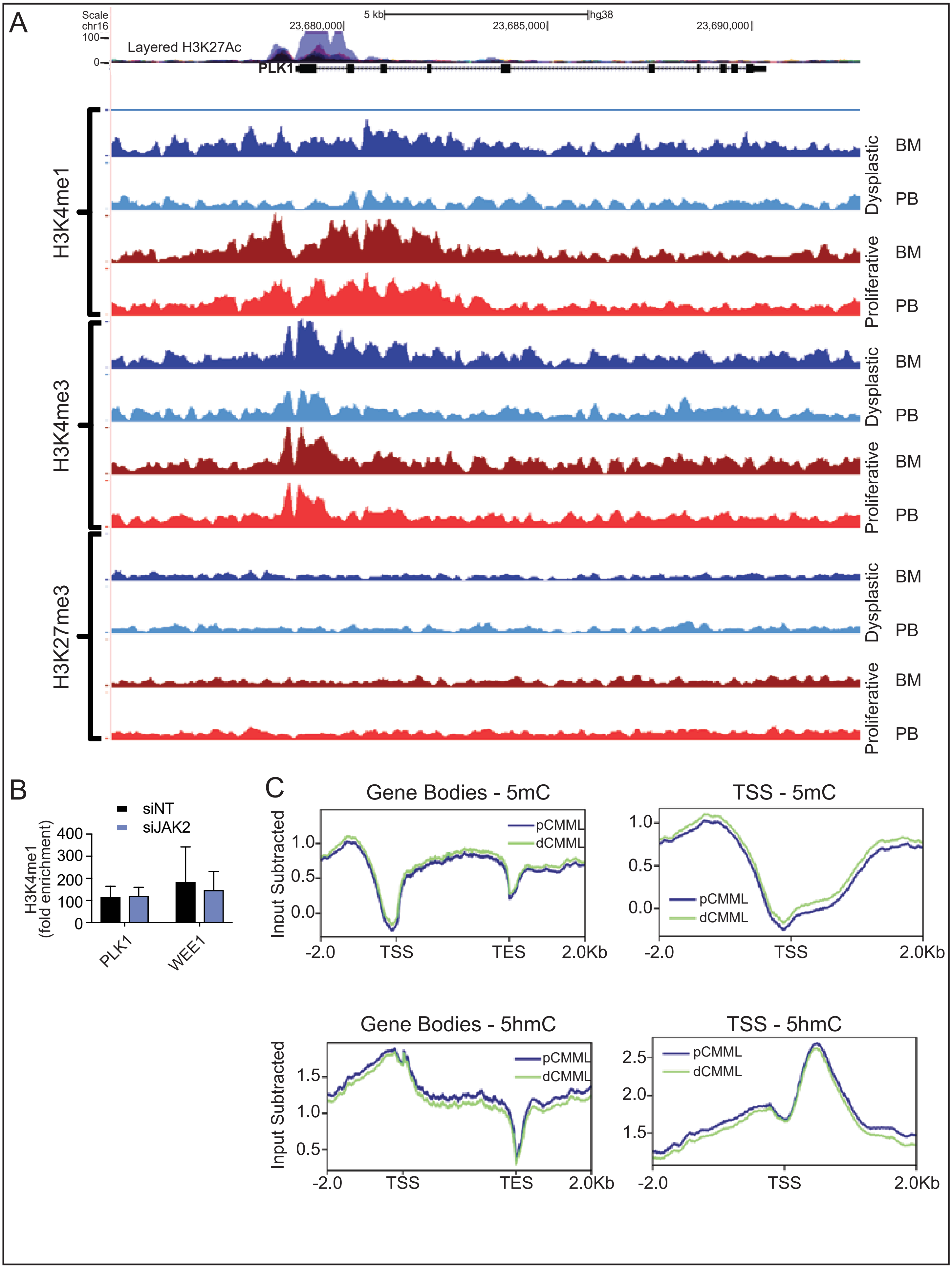

**Supplementary Figure 6**

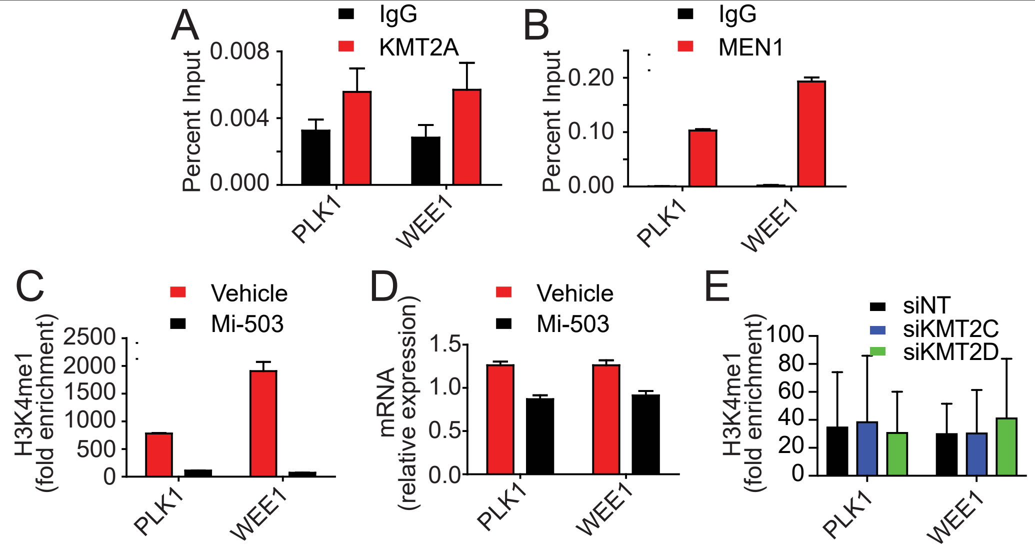

### Supplementary Figure 7

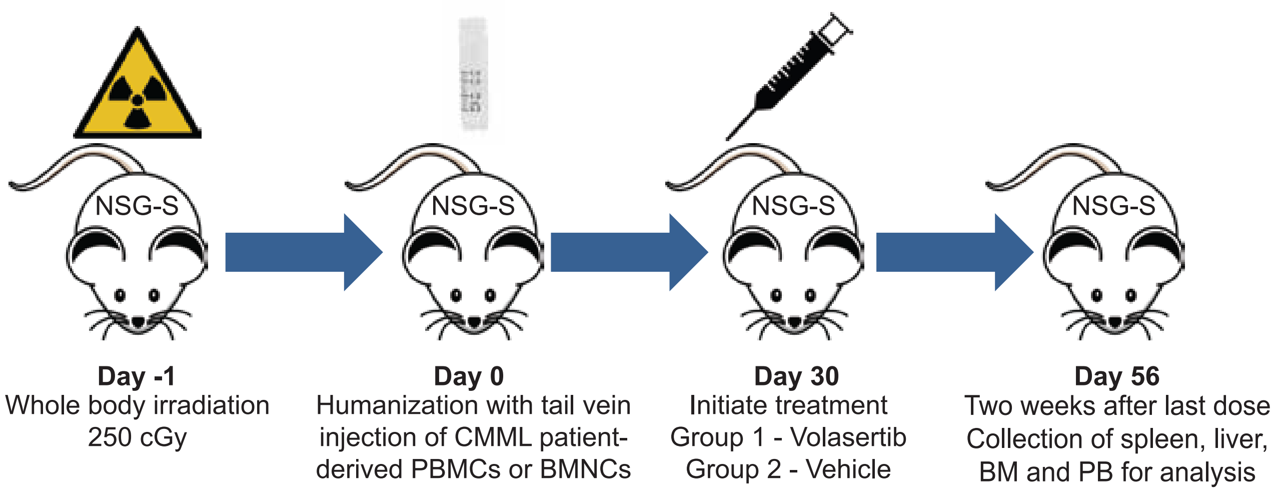

**Figure S1. RAS pathway mutations are associated with poor outcomes and are enriched in pCMML A.** Kaplan-Meier curve depicting overall survival in CMML stratified based on RAS pathway mutation status. **B.** Kaplan-Meier curve depicting overall survival of CMML patients stratified by *NRAS* mutation status. **C.** Kaplan-Meier curve depicting AML free survival of CMML patients stratified by *NRAS* mutation status. **D.** Violin plots representing variant allele frequencies (VAFs) of *NRAS*, *KRAS* and *CBL* mutations in dCMML and pCMML. * indicates p value < 0.05.

**Figure S2. Driver mutation alterations encountered in CMML progression to AML A.** Driver mutations and somatic copy number alterations (SCNAs) in CMML patients that progressed to sAML. For each patient (individual columns) putative driver events at CMML (blue) and sAML (red) are depicted. Clinical characteristics correspond to measurements at the CMML stage. **B.** Categorization of mutations detected in CMML and sAML sample pairs. 1d are mutations detected in both CMML and sAML and considered as primary drivers, 2d are those detected only in sAML and considered as secondary drivers, and, 3d are those detected in CMML and lost in sAML, suggesting subclonal secondary drivers. **C.** Prevalence of the main driver mutations and SCNAs in the categories of primary driver (x-axis) events (1d) and secondary driver events (2d), occurring during leukemic transformation (LT) (y-axis). Confidence intervals represent Fisher exact test alpha = 0.05 for point mutations. **D.** Fraction of *NRAS* mutations among all drivers in CMML (1d), gained (2d) or lost (3d) during LT. **E.** Diagrams of CMML with (below) and without (above) RAS-driven subclones. CMML without *RAS* mutations are at increased risk for acquisition of *RAS* mutations during LT. * indicates p value < 0.05. **F.** Kaplan-Meier curve depicting overall survival of CMML with and without *RAS* mutations during LT.

**Figure S4. Mitotic kinase enrichment in pCMML is likely unrelated to bulk sequencing or *JAK2* mutation. A.** Pearson's correlation comparing RNA-seq data presented as median log2-normalized transcript number. Measured transcripts from unsorted PB MNCs (n=5) were compared to transcripts from CD34+ sorted progenitor cells (left, n=5) as well as CD14+ sorted monocytes (right, n=5). **B.** qPCR assessing relative expression of PLK1 (left) and WEE1 (right) in pCMML cases with a *JAK2* mutation (n=4) and those with a *NRAS* mutation (n=21). Data presented as mean ± SEM.

**Figure S5. Sequence-specific analyses reveal no differences in common epigenetic marks at *PLK1* and *WEE1* loci between dCMML and pCMML. A.** Representative sequence-specific ChIP-seq data demonstrating relative H3K4me1, H3K4me3 and H3K27me3 signals comparing dCMML patient-derived BM (dark blue) and PB (light blue) and pCMML patient-derived BM (dark red) and PB (light red). Data is depicted on the PLK1 gene map above. Blue and purple layered traces are H3K27Ac occupancy data from UCSC Genome browser. **B.** Tag density plots from DNA immunoprecipitation sequencing (DIP-seq) assessing differences in 5-methylcytosine (5-mC, top) and 5-hydroxymethylcytosine (5-hmC, bottom) between BM-derived MNCs from pCMML (n=18) and dCMML (n=9). Comparisons depicted include differences in global gene bodies (left) and at transcriptional start sites (TSS) of *PLK1* and *WEE1* (right).

**Figure S6. KMT2A and not KMT2C/D regulate PLK1 and WEE1 expression in pCMML. A and B.** ChIP-PCR assessing enrichment of KMT2A (A) and MEN1 (B) at promoters of *PLK1* and *WEE1* relative to isotype control (IgG) in *NRAS* mutant pCMML patient-derived MNCs. **E and F.** ChIP-PCR assessing H3K4me1 enrichment at promoters of *PLK1* and *WEE1* (E) and qPCR assessing PLK1 and WEE1 levels (F) in *NRAS* mutant CMML patient-derived MNCs after treatment with either vehicle control or Mi-503. Data are presented as mean ± SEM from three individual patient-derived samples.

**Figure S7. Patient-derived xenograft drug study design.** Diagram depicting the general experimental design of PDX drug studies.

**Table S1. Clinical and laboratory features and subsequent events in 1183 WHO defined patients with chronic myelomonocytic leukemia (CMML) stratified by Mayo Clinic, Austrian and French (GFM) cohorts, Related to Figure 1**

| *Variables [Median or n; range or %]* | **All patients (*n=1183* )** | **Mayo Clinic Cohort (*n*=397)** | **Austrian Cohort (*n*=175)** | **GFM Cohort (n=417)** | **P value** |
|---|---|---|---|---|---|
| **Age in years; median (range) Evaluable= 1181** | 72 (18.1-95.2) | 70.6 (18.1-95.2) | 72 (45-93) | 73.64 (28.6-92.97) | **0.0001** |
| **Sex (Male); *n* (%) Evaluable= 1165** | 774 (66) | 397 (67) | 106 (61) | 271 (68) | 0.21 |
| **Hemoglobin g/dL; median (range) Evaluable= 1162** | 11.1 (4.2-18) | 10.7 (4.3-17) | 11 (5.8-15.3) | 11.7 (4.2-17.7) | **0.0001** |
| **WBC x $10^9$/L; median (range) Evaluable= 1183** | 12.6 (1.3-366.8) | 12.9 (1.3-264.8) | 17 (2.5-156) | 11.3 (1.9-366.8) | **0.0013** |
| **ANC x$10^9$/L; median (range) Evaluable= 576** | 6.2 (0-151) | 6.2 (0-151) | NA | NA | **-** |
| **AMC x $10^9$/L; median (range) Evaluable= 1152** | 2.7 (1-102.5) | 3 (1-84) | 3.62 (1.015-54.4) | 2.2 (1-102.5) | **0.1** |
| **Platelets x $10^9$/L; median(range) Evaluable= 1162** | 108 (3-1427) | 101 (7-1277) | 108 (5-726) | 123 (3-1427) | **0.0002** |
| **IMC (Y/N) Evaluable= 960** | 546 (57) | 348 (60) | 40 (73) | 158 (49) | **0.0005** |
| **PB blasts %; median (range) Evaluable= 1056** | 0 (0-19) | 0 (0-19) | 0 (0-17) | 0 (0-14) | **0.0002** |
| **BM blasts %; median (range) Evaluable= 941** | 4 (0-19) | 3 (0-19) | NA | 5 (0-19) | **-** |
| **FAB CMML diagnosis Evaluable= 1183** | | | | | |
| **dCMML; *n* (%)** | 607 (51) | 298 (50) | 72 (41) | 237 (57) | **0.0019** |
| **pCMML; *n* (%)** | 576 (49) | 293 (50) | 103 (59) | 180 (43) | |
| **WHO 2016 CMML diagnosis Evaluable= 1065** | | | | | |
| **CMML-0; n (%)** | 530 (50) | 323 (55) | 75 (45) | 132 (43) | **0.0002** |
| **CMML-1; n (%)** | 304 (29) | 158 (27) | 41 (24) | 105 (34) | |
| **CMML-2; n (%)** | 231 (22) | 106 (18) | 52 (31) | 73 (24) | |
| **Mayo-French cytogenetic risk stratification; *n* (%) Evaluable= 1032** | | | | | |

| | | | | | |
|---|---|---|---|---|---|
| **Low** | 772 (75) | 417 (74) | 77 (66) | 278 (79) | 0.03 |
| **Intermediate** | 208 (20) | 111 (20) | 31 (27) | 66 (19) | |
| **High** | 52 (5) | 34 (6) | 8 (7) | 10 (3) | |
| **Next generation sequencing analysis; n (%) Evaluable=977** | | | | | |
| **1. Epigenetic regulators** | | | | | |
| ***TET2*** | 517 (53) | 193 (50) | 80 (46) | 244 (60) | **0.0015** |
| ***IDH1*** | 9 (1) | 5 (1) | NA | 4 (1) | - |
| ***IDH2*** | 41 (5) | 20 (5) | NA | 21 (5) | - |
| ***DNMT3A*** | 49 (6) | 16 (4) | 11 (6) | 22 (7) | 0.25 |
| **2. Chromatin regulators** | | | | | |
| ***ASXL1*** | 365 (37) | 187 (48) | 45 (26) | 133 (32) | **<0.0001** |
| ***EZH2*** | 54 (6) | 16 (4) | 18 (10) | 20 (6) | 0.017 |
| **3. Transcription factors** | | | | | |
| ***RUNX1*** | 76 (14) | 49 (13) | 27 (15) | NA | - |
| **4. Spliceosome factors** | | | | | |
| ***SRSF2*** | 412 (44) | 177 (46) | 69 (39) | 166 (44) | 0.39 |
| ***SF3B1*** | 58 (6) | 17 (4) | 14 (8) | 27 (7) | 0.15 |
| ***U2AF1*** | 51 (7) | 27 (7) | NA | 24 (6) | - |
| ***ZRSR2*** | 38 (5) | 16 (4) | NA | 22 (6) | - |
| **5. Cell signaling** | | | | | |
| ***NRAS*** | 149 (15) | 67 (17) | 32 (18) | 50 (12) | 0.055 |
| ***KRAS*** | 87 (9) | 20 (5) | 16 (9) | 51 (12) | **0.002** |
| ***CBL*** | 145 (15) | 67 (17) | 28 (16) | 50 (12) | 0.10 |
| ***PTPN11*** | 26 (5) | 14 (4) | 12 (7) | NA | - |
| ***JAK2*** | 73 (7) | 25 (6) | 17 (10) | 31 (8) | 0.39 |
| ***CSF3R*** | 29 (4) | 4 (1) | NA | 25 (7) | **-** |
| ***KIT*** | 18 (3) | 13 (3) | NA | 5 (2) | 0.13 |
| ***MPL*** | 3 (1) | 3 (1) | NA | NA | - |
| ***CALR*** | 0 | 0 | NA | NA | - |
| **6. Tumor suppressor gene** | | | | | |
| ***Tp53*** | 25 (3) | 12 (3) | 8 (5) | 5 (2) | 0.42 |
| **7. Others** | | | | | |
| ***SETBP1*** | 69 (9) | 40 (10) | 15 (9) | 14 (8) | 0.56 |
| **Mayo Molecular Model Evaluable= 743** | | | | | |
| **Low risk; n (%)** | 89 (12) | 32 (8) | 9 (17) | 48 (15) | **<0.0001** |
| **Intermediate-1 risk; n (%)** | 230 (31) | 95 (25) | 14 (26) | 121 (39) | |
| **Intermediate-2 risk; n (%)** | 231 (31) | 131 (35) | 14 (26) | 86 (27) | |
| **High risk; n (%)** | 193 (26) | 119 (32) | 16 (30) | 58 (19) | |
| **GFM Prognostic Model Evaluable= 938** | | | | | |
| **Low risk; n (%)** | 420 (45) | 154 (40) | 65 (37) | 201 (53) | |

| | | | | | |
|---|---|---|---|---|---|
| **Intermediate risk; n (%)** | 359 (38) | 145 (40) | 83 (48) | 131 (34) | **<0.0001** |
| **High risk; n (%)** | 159 (17) | 85 (22) | 26 (15) | 48 (13) | |
| **CPSS-Mol Prognostic Model Evaluable= 339** | | | | | |
| **Low risk; n (%)** | 38 (11) | 38 (11) | NA | NA | - |
| **Intermediate-1 risk; n (%)** | 86 (25) | 86 (25) | NA | NA | |
| **Intermediate-2 risk; n (%)** | 131 (39) | 131 (39) | NA | NA | |
| **High risk; n (%)** | 84 (25) | 84 (25) | NA | NA | |
| **Deaths; *n* (%) Evaluable= 995** | 652 (55) | 394 (67) | 112 (64) | 146 (35) | **<0.0001** |
| **Leukemic transformation; n (%) Evaluable= 995** | 196 (20) | 119 (20) | 12 (7) | 65 (28) | **<0.0001** |

**The bold values represent p values that are statistically significant; *p* < 0.05 (Only provided if data is available for all three cohorts).**

**Key:** CMML: chronic myelomonocytic leukemia, AMC: absolute monocyte count; ANC: absolute neutrophil count; IMC: immature circulating cells; WBC: white blood cell count; PB: peripheral blood; BM: bone marrow; WHO: World Health Organization; CPSS-Mol: clinical/molecular CMML-specific prognostic scoring system; dCMML: dysplastic chronic myelomonocytic leukemia; pCMML: proliferative chronic myelomonocytic leukemia; FAB: French-American-British; GFM: Groupe Francophone des Myelodyplasies; IMC: immature myeloid cells; $^{MT}$: mutated; $^{WT}$: wild type; CI: confidence interval ; NA: not available.

**Table S2. Clinical, pathological and molecular characteristics of 1183 WHO defined chronic myelomonocytic leukemia (CMML) patients stratified by proliferative and dysplastic subtypes at diagnosis, Related to Figure 1**

| *Variables [Median or n; range or %]* | **All patients (*n=1183* )** | **pCMML (n=576)** | **dCMML (n=607)** | **P value** |
|---|---|---|---|---|
| **Age in years; median (range) Evaluable= 1181** | 72 (18.1-95.2) | 71.34 (18.1-93) | 73 (28.3-95.1) | **0.004** |
| **Sex (Male); *n* (%) Evaluable= 1165** | 774 (66) | 35 (63) | 419 (70) | **0.02** |
| **Hemoglobin g/dL; median (range) Evaluable= 1162** | 11.1 (4.2-18) | 10.9 (4.2-17.7) | 11.2 (4.6-16.8) | **0.01** |
| **WBC x $10^9$/L; median (range) Evaluable= 1183** | 12.6 (1.3-366.8) | 24.1 (13-366.8) | 7.2 (1.3-12.9) | **<0.0001** |
| **ANC x$10^9$/L; median (range) Evaluable= 576** | 6.2 (0-151) | 13.55 (1.5-151) | 3.1 (0-11) | **<0.0001** |
| **AMC x $10^9$/L; median (range) Evaluable= 1152** | 2.5 (1-102.5) | 4.7 (1-102.5) | 1.5 (1-27.3) | **<0.0001** |
| **Platelets x $10^9$/L; median(range) Evaluable= 1162** | 108 (3-1427) | 116 (5-1427) | 104 (3-1051) | 0.19 |
| **IMC (Y/N) Evaluable= 960** | 546 (57) | 340 (74) | 206 (41) | **<0.0001** |
| **PB blasts %; median (range) Evaluable= 1056** | 0 (0-19) | 0 (0-19) | 0 (0-17) | **<0.0001** |
| **BM blasts %; median (range) Evaluable= 941** | 4 (0-19) | 4 (0-19) | 4 (0-19) | 0.41 |
| **WHO 2016 CMML diagnosis Evaluable= 1065** | | | | |
| **CMML-0; n (%)** | 530 (50) | 240 (46) | 290 (54) | **0.03** |
| **CMML-1; n (%)** | 304 (29) | 159 (30) | 145 (27) | |
| **CMML-2; n (%)** | 231 (22) | 126 (24) | 105 (19) | |
| **Mayo-French cytogenetic risk stratification; *n* (%) Evaluable= 1032** | | | | |
| **Low** | 772 (75) | 361 (72) | 411 (77) | 0.09 |
| **Intermediate** | 208 (20) | 115 (23) | 93 (18) | |
| **High** | 52 (5) | 25 (5) | 27 (5) | |

| | | | | |
|---|---|---|---|---|
| **Next generation sequencing analysis; n (%) Evaluable=977** | | | | |
| **1. Epigenetic regulators** | | | | |
| ***TET2*** | 517 (53) | 227 (48) | 290 (58) | **0.0022** |
| ***IDH1*** | 9 (1) | 3 (1) | 6 (1) | 0.44 |
| ***IDH2*** | 41 (5) | 18 (5) | 23 (5) | 0.78 |
| ***DNMT3A*** | 49 (6) | 24 (6) | 25 (6) | 0.93 |
| **2. Chromatin regulators** | | | | |
| ***ASXL1*** | 365 (37) | 221 (47) | 144 (29) | **<0.0001** |
| ***EZH2*** | 54 (6) | 38 (9) | 16 (3) | **0.0011** |
| **3. Transcription factors** | | | | |
| ***RUNX1*** | 76 (14) | 36 (12) | 40 (15) | 0.36 |
| **4. Spliceosome factors** | | | | |
| ***SRSF2*** | 412 (44) | 202 (44) | 210 (43) | 0.8 |
| ***SF3B1*** | 58 (6) | 22 (5) | 36 (7) | 0.1 |
| ***U2AF1*** | 51 (7) | 25 (7) | 26 (6) | 0.63 |
| ***ZRSR2*** | 38 (5) | 9 (3) | 29 (7) | **0.0041** |
| ***5.* Cell signaling** | | | | |
| ***NRAS*** | 149 (15) | 110 (23) | 39 (8) | **<0.0001** |
| ***KRAS*** | 87 (9) | 43 (9) | 44 (9) | 0.84 |
| ***CBL*** | 145 (15) | 85 (18) | 60 (12) | **0.009** |
| ***PTPN11*** | 26 (5) | 14 (5) | 12 (5) | 0.9 |
| ***JAK2*** | 73 (7) | 51 (11) | 22 (4) | **0.0002** |
| ***CSF3R*** | 29 (4) | 15 (4) | 14 (4) | 0.58 |
| ***KIT*** | 18 (3) | 11 (3) | 7 (2) | 0.24 |
| ***MPL*** | 3 (1) | 2 (1) | 1 (1) | 0.55 |
| ***CALR*** | 0 | 0 | 0 | **-** |
| **6. Tumor suppressor gene** | | | | |
| ***Tp53*** | 25 (3) | 12 (3) | 13 (3) | 0.78 |
| **7. Others** | | | | |
| ***SETBP1*** | 69 (9) | 42 (11) | 27 (8) | 0.1 |
| **Mayo Molecular Model Evaluable= 743** | | | | |
| **Low risk; n (%)** | 89 (12) | 23 (7) | 66 (17) | **<0.0001** |
| **Intermediate-1 risk; n (%)** | 230 (31) | 82 (23) | 148 (38) | |
| **Intermediate-2 risk; n (%)** | 231 (31) | 113 (32) | 118 (30) | |
| **High risk; n (%)** | 193 (26) | 135 (38) | 58 (15) | |
| **GFM Prognostic Model Evaluable= 938** | | | | |
| **Low risk; n (%)** | 420 (45) | 62 (14) | 358 (74) | **<0.0001** |
| **Intermediate risk; n (%)** | 359 (38) | 255 (56) | 104 (21) | |
| **High risk; n (%)** | 159 (17) | 135 (30) | 24 (5) | |
| **CPSS-Mol Prognostic Model Evaluable= 339** | | | | |

| | | | | |
|---|---|---|---|---|
| **Low risk; n (%)** | 38 (11) | 0 | 38 (22) | **<0.0001** |
| **Intermediate-1 risk; n (%)** | 86 (25) | 25 (15) | 61 (35) | |
| **Intermediate-2 risk; n (%)** | 131 (39) | 76 (46) | 55 (32) | |
| **High risk; n (%)** | 84 (25) | 65 (39) | 19 (11) | |
| **Deaths; *n* (%) Evaluable= 995** | 652 (55) | 346 (60) | 306 (50) | **0.0008** |
| **Leukemic transformation; n (%) Evaluable= 995** | 196 (20) | 99 (20) | 97 (19) | 0.53 |

**The bold values represent p values that are statistically significant; *p* < 0.05.**

**Key:** CMML: chronic myelomonocytic leukemia, AMC: absolute monocyte count; ANC: absolute neutrophil count; IMC: immature circulating cells; WBC: white blood cell count; PB: peripheral blood; BM: bone marrow; WHO: World Health Organization; CPSS-Mol: clinical/molecular CMML-specific prognostic scoring system; dCMML: dysplastic chronic myelomonocytic leukemia; pCMML: proliferative chronic myelomonocytic leukemia; FAB: French-American-British; GFM: Groupe Francophone des Myelodyplasies; IMC: immature myeloid cells; $^{MT}$: mutated; $^{WT}$: wild type; CI: confidence interval

**Table S3. Clinical and pathological characteristics of CMML patients tested through a research-based whole exome sequencing (WES) platform, Related to Figure 2**

| ***Variables [Median or n; range or %]*** | ***CMML WES cohort (n=48)*** |
|---|---|
| **Age in years; median (range)** | 69.55 (18.04-86.82) |
| **Sex (Male); *n* (%)** | 28 (58) |
| **Hemoglobin g/dL; median (range)** | 10.85 (4.3-16) |
| **WBC x $10^9$/L; median (range)** | 13.8 (2.8-264.8) |
| **ANC x$10^9$/L; median (range)** | 83.25 (0.84-37839.92) |
| **AMC x $10^9$/L; median (range)** | 2.8 (1-24.8) |
| **Platelets x $10^9$/L; median (range)** | 98.5 (14-362) |
| **IMC (Y/N) *n* (%)** | 30 (63) |
| **PB blasts %; median (range)** | 0 (0-19) |
| **BM blasts %; median (range)** | 4 (0-18) |
| **FAB CMML diagnosis** | |
| **dCMML; *n* (%)** | 22 (46) |
| **pCMML; *n* (%)** | 26 (54) |
| **WHO 2016 CMML diagnosis** | |
| **CMML-0; n (%)** | 18 (38) |
| **CMML-1; n (%)** | 17 (35) |
| **CMML-2; n (%)** | 13 (27) |
| **Mayo-French cytogenetic risk stratification; *n* (%)** | |
| **Low** | 27 (56) |
| **Intermediate** | 9 (19) |
| **High** | 6 (13) |
| **Mayo Molecular Model** | |
| Low risk; n (%) | 3 (63) |
| Intermediate-1 risk; n (%) | 10 (21) |
| Intermediate-2 risk; n (%) | 4 (8) |
| High risk; n (%) | 11 (23) |
| **GFM Prognostic Model** | |

| | |
|---|---|
| Low risk; n (%) | 10 (21) |
| Intermediate risk; n (%) | 12 (25) |
| High risk; n (%) | 7 (15) |
| **CPSS-Mol Prognostic Model** | |
| Low risk; n (%) | 3 (63) |
| Intermediate-1 risk; n (%) | 8 (17) |
| Intermediate-2 risk; n (%) | 3 (63) |
| High risk; n (%) | 12 (25) |
| **Deaths; *n* (%)** | 38 (80) |
| **Leukemic transformation; n (%)** | 48 (100) |

**Key:** CMML: chronic myelomonocytic leukemia, WES: whole exome sequencing, AMC: absolute monocyte count; ANC: absolute neutrophil count; IMC: immature circulating cells; WBC: white blood cell count; PB: peripheral blood; BM: bone marrow; WHO: World Health Organization; CPSS-Mol: clinical/molecular CMML-specific prognostic scoring system; dCMML: dysplastic chronic myelomonocytic leukemia; pCMML: proliferative chronic myelomonocytic leukemia; FAB: French-American-British; GFM: Groupe Francophone des Myelodyplasies; IMC: immature myeloid cells; $^{MT}$: mutated; $^{WT}$: wild type; CI: confidence interval
